# Therapeutic mitigation of measles-like immune amnesia and exacerbated disease after prior respiratory virus infections in ferrets

**DOI:** 10.1101/2023.09.01.555992

**Authors:** Robert M Cox, Josef D Wolf, Nicole A Lieberman, Carolin M Lieber, Hae-Ji Kang, Zachary M Sticher, Jeong-Joong Yoon, Meghan K Andrews, Mugunthan Govindarajan, Rebecca E Krueger, Elizabeth B. Sobolik, Michael G Natchus, Andrew T Gewirtz, Rik deSwart, Alexander A Kolykhalov, Khan Hekmatyar, Kaori Sakamoto, Alexander L Greninger, Richard K Plemper

## Abstract

After years of the COVID-19 pandemic, over 40 million children worldwide are at risk of measles due to delayed vaccination^1^ and temporary SARS-CoV-2 viral dominance^2^. Acute measles has a case-fatality rate of ∼1%, but most morbidity and mortality arise post-measles due to destruction of pre-existing immune memory by lymphotropic measles virus (MeV)^3,4^, a paramyxovirus of the *Morbillivirus* genus. MeV-induced immune amnesia is not mitigated by post-exposure vaccination and the impact of unrelated respiratory virus disease history on measles severity has not been defined. We used a lethal canine distemper virus (CDV)-ferret model as surrogate for human morbillivirus disease^5^ and employed the orally efficacious broad-spectrum paramyxovirus polymerase inhibitor GHP-88309^6^ to establish measles treatment paradigms. Applying a receptor tropism-intact recombinant CDV with low lethality, we provide *in vivo* confirmation of the morbillivirus immune amnesia hypothesis and reveal an 8-day advantage of antiviral treatment versus therapeutic vaccination in preserving immune memory. Infection of ferrets with non-lethal influenza A virus (IAV) A/CA/07/2009 (H1N1) or respiratory syncytial virus (RSV) four weeks prior to CDV caused exacerbated CDV disease that rapidly advanced to fatal hemorrhagic pneumonia associated with lung onslaught by commensal bacteria. RNAseq of BAL samples and lung tissue identified CDV-induced expression of trefoil factor (TFF) peptides, which was absent in animals pre-infected with IAV, thus highlighting that immune priming by unrelated respiratory viruses influences morbillivirus infection outcome. Non-invasive pulmonary ferret MRI revealed that severe outcomes of consecutive IAV/CDV infections were prevented by oral GHP-88309 treatment even when initiated after peak clinical signs of CDV. These findings validate the morbillivirus immune amnesia hypothesis, define treatment paradigms for measles, identify prior disease history as risk factor for exacerbated morbillivirus disease, and demonstrate that treating morbillivirus infection with direct-acting oral antivirals provides therapeutic benefit regardless of whether the time window to mitigate primary clinical signs of infection has closed.

## Main Text

Morbilliviruses such as MeV and CDV invade their hosts through infection of alveolar macrophages (AMs) and dendritic cells (DCs), using signaling lymphocyte activation molecule (CD150) as receptor^7^. Subsequently, peripheral blood mononuclear cell (PBMC)-supported cell-associated viremia ensues, until viral re-entry into the respiratory tract and infection of epithelial cells through the basolaterally-expressed Nectin-4 receptor^8^. Viral replication during the viremic phase in CD150^+^ lymphocytes in peripheral blood and primary, secondary, and tertiary lymphoid tissues depletes CD150-positive lymphocytes, including memory T and B cells, reducing host antibody repertoire and erasing immune memory^3,4^. Such MeV-induced immune amnesia is hypothesized to increase vulnerability to opportunistic infections, leading to significantly increased morbidity and mortality rates from unrelated infectious diseases in the years following primary measles^9^. Treatment options of acute measles are limited to supportive care, IgG therapy in some high-income countries^10^, and quarantine of the patient. The exceptionally high infectivity of MeV^11^ and vaccine hesitancy resulted in a measles resurgence in many European countries in 2019 even prior to the COVID-19 pandemic^12^. Yet temporary pausing of vaccination campaigns in low and middle-income countries during the pandemic has exacerbated the MeV resurgence and major measles outbreaks are anticipated globally^12^. Despite the severity of this health threat, experimental insight is lacking into the time window for prevention of MeV-induced immune-suppression through direct-acting antivirals (DAAs) and the consequences of a large measles wave coinciding with high activity of unrelated respiratory viruses such as influenza virus and RSV.

We previously developed an orally efficacious morbillivirus polymerase inhibitor, ERDRP-0519, which fully protects ferrets against a lethal CDV infection in a post-exposure prophylactic dosing regimen^13^. We recently identified a structurally and mechanistically distinct broadened-spectrum paramyxovirus polymerase inhibitor, GHP-88309, which is orally efficacious against parainfluenzaviruses and blocks morbilliviruses with sub-micromolar potency in cell culture^6^. Using these compounds, we established in this work measles treatment paradigms and explored the effect of prior disease history on severity of morbillivirus infection.

### GHP-88309 is orally bioavailable in ferrets

To verify efficient oral delivery of GHP-88309 to ferrets, we determined single-dose pharmacokinetic (PK) profiles after oral administration at 50 and 150 mg/kg bodyweight. GHP-88309 demonstrated dose-dependent plasma exposure of 177.8 hrs×nmol/ml and 754.1 hrs×nmol/ml, respectively, and sustained tissue distribution exceeding 1 nmol/g tissue 12 hours after administration (Extended Data Fig. 1a,b; Supplementary Table S1). Twice daily (b.i.d.) oral dosing at 15 and 50 mg/kg in a 14-day non-formal tolerability study revealed no signs of drug-induced toxicity or abnormalities in serum chemistry (Supplementary Fig. S1a-d) and an abbreviated repeated-dose PK study confirmed consistent exposure on day 7 (Supplementary Fig. S2; Supplementary Table S2). Average trough GHP-88309 plasma concentration in the 50 mg/kg b.i.d. group was 2.4 µM, which was equivalent to >2 × EC_90_ against CDV^6^. This dose of GHP-88309 was used for all subsequent experiments.

### Oral GHP-88309 expands to 8-days the time window of full protection compared to near-exposure vaccination

Efficacy of GHP-88309, ERDRP-0519, and therapeutic vaccination was compared in CDV-naïve ferrets infected with reporter-free, highly pathogenic recombinant recCDV-5804p (recCDV)^14^, followed by oral treatment with GHP-88309 (50 mg/kg, b.i.d.) initiated 3, 5, or 7 days post infection (dpi) or oral ERDRP-0519 (50 mg/kg, b.i.d) started 3 dpi (Fig. 1a). Animals in vaccination groups received Purevax CDV vaccine 28, 3, or 1 day prior to, or 1 day after, infection in a prime-boost (28-day group; –28-day prime, –14-day boost) or prime only (all other groups) regimen. A reference group received CDV vaccine at 2 months of age. All animals were fully vaccinated against rabies virus (RABV) by their supplier and confirmed to have α-RABV antibodies prior to use (Supplementary Fig. S3). Ferrets were monitored daily for clinical signs of morbillivirus disease (fever, rash, loss of bodyweight) and in regular intervals for PBMC-associated viremia, complete blood counts (CBC), and titer of α-RABV neutralizing antibodies (nAbs) (Fig. 1a). All vehicle-treated animals succumbed within 12 days of infection (Fig. 1b) and developed severe clinical signs (Extended Data Fig. 2a-e), whereas GHP-88309 mediated complete survival and statistically significantly reduced virus load (Fig. 1c) when treatment was initiated 3 or 5 dpi, at the onset of viremia and rash, respectively. Treatment started 7 dpi did not significantly improve outcome, and ERDRP-0519 treatment started 3 dpi only partially protected against infection with this wild type recCDV, which is more virulent than the fluorescent recCDV reporter strain used previously^13^. Fully prime-boost vaccinated animals were protected, but Neither post-exposure nor 1-day pre-exposure vaccination was efficacious and, moreover, 2 of 3 animals vaccinated prophylactically 3 days prior to infection also succumbed. Clinical signs in animals of all GHP-88309 treatment groups, except the 7-dpi arm, were unremarkable 12 dpi (Fig. 1d; Extended Data Fig. 2), whereas ERDRP-0519-treated animals showed moderate, and vehicle-treated animals severe, clinical signs.

**Figure 1.**
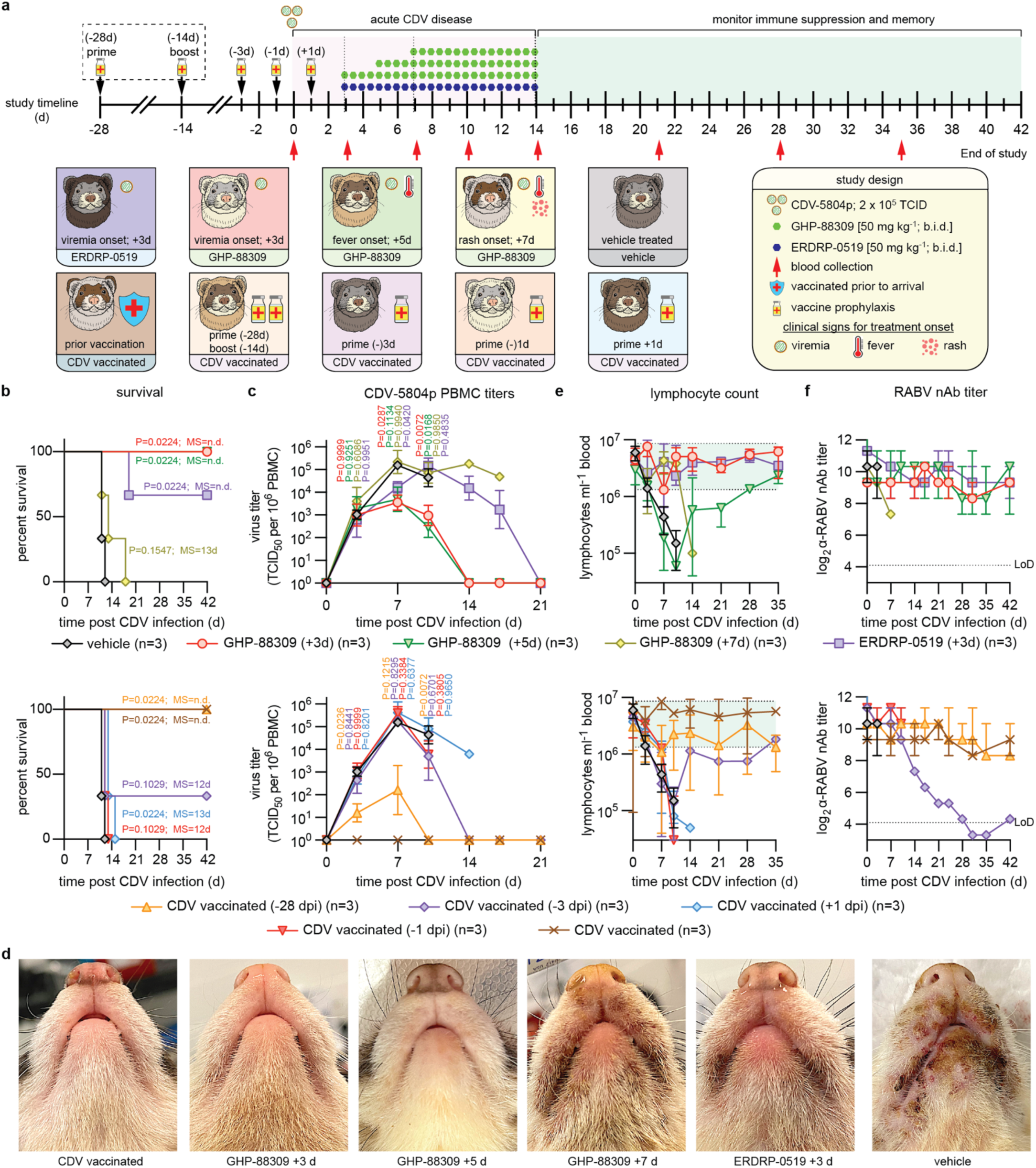
Therapeutic treatment of lethal CDV infection in ferrets. Ferrets were infected with a lethal challenge of CDV and treated with GHP-88309, ERDRP-0519, or therapeutic vaccination. **a**, Schematic of the study design. Ferrets were infected with wild-type recCDV-5804p and monitored for 6 weeks. **b**, Survival curves of ferrets infected in (a). log-rank (Mantel-Cox) test, median survival is stated. **c**, PBMC associated viremia titers of CDV infected ferrets shown in (a). 2-way ANOVA with Tukey’s post-hoc test. **d**, Images of ferrets taken 12 days after infection with recCDV-5804p. **e**, Lymphocyte counts from infected ferrets measured during the duration of the study detailed in (a). Green shading denotes normal range. **f**, RABV neutralizing antibody titers from CDV infected ferrets. Symbols (c, e, f) represent geometric means ± geometric SD, lines intersect means. In (b, c, d, e), top row shows results for inhibitor-treated groups, bottom row for vaccinated animals; LoD, limit of detection; n=3.

Lymphocyte counts in the vehicle group dropped rapidly by approximately two orders of magnitude by 12 dpi (Fig. 1e). Both GHP-88309 and ERDRP-0519 treatment initiated up to 3 dpi prevented lymphocytopenia. GHP-88309 started 5 dpi did not alleviate initial PBMC collapse, but resulted in populations rapidly recovering such that lymphocytopenia resolved within 4 weeks. Fully vaccinated animals did not develop lymphocytopenia. However, the single surviving animal of the –3 dpi prophylactic vaccination group also experienced temporary collapse of the PBMC population. Pre-existing unrelated humoral immunity, assessed using α-RABV nAbs as a biomarker, was fully preserved in animals of the 5 dpi and earlier treatment groups, but was permanently destroyed in the surviving animal of the –3 dpi vaccination group (Fig. 1f). Thus, GHP-88309 prevents acute lethal morbillivirus infection and mitigates lymphopenia up to 5 days post-infection thereby providing extending the intervention opportunity by at least 8 days compared to near-exposure vaccination. Based on superior oral efficacy of GHP-88309 compared to ERDRP-0519, we selected GHP-88309 for all subsequent experiments.

### Confirmation of immune amnesia hypothesis via recCDV with tunable polymerase processivity

The 100% lethality of recCDV in untreated ferrets within 2 weeks of exposure precluded use of this model in exploring questions relevant to human morbillivirus-induced disease in which lethality is slower and less uniform. To surmount this hurdle, we engineered an attenuated recCDV with intact viral receptor tropism. Previously, we demonstrated that variable length deletions of 39-55 amino acids in the structurally disordered CDV nucleocapsid (N) protein tail domain affect polymerase processivity to different degrees, resulting in genetically stable tunable recCDV attenuation^15^. Employing a 55-amino acid deletion recCDV Ný425-479 strain, we compared virulence and severity of immune suppression with that of unmodified recCDV and a previously described tropism-altered recCDV Nectin-4-blind mutant strain^3^ that cannot engage the Nectin-4 epithelial cell receptor (Fig. 2a).

**Figure 2.**
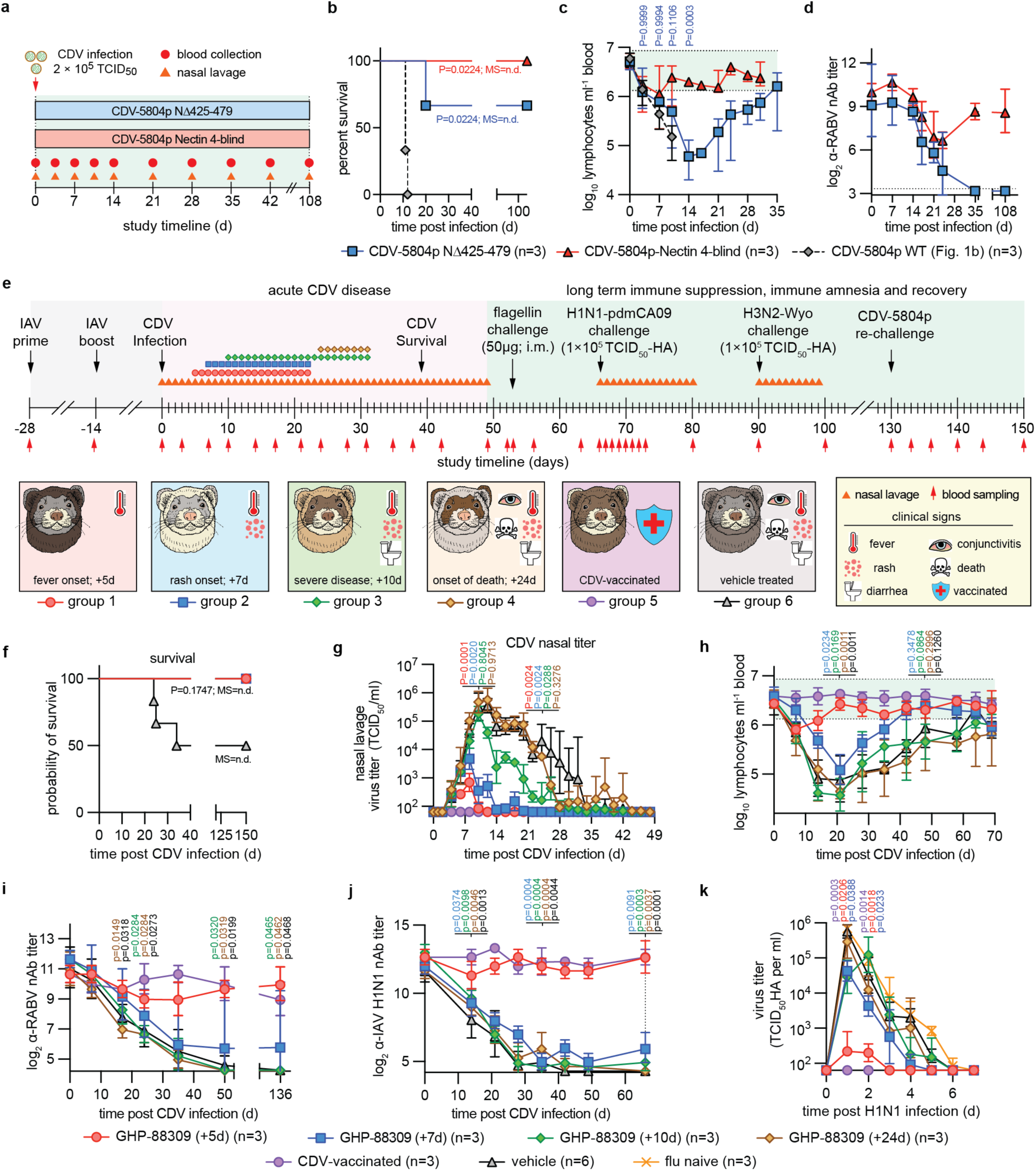
Ferret model of CDV-induced immune amnesia. a, Study schematic. Ferrets were infected intranasally with recCDV-5804p NΔ425-479 or recCDV-5804p-Nectin 4-blind and monitored for 108 days. Nasal lavages were taken from recCDV-5804p NΔ425-479 infected ferrets only. Blood was sampled regularly from all animals. **b-d,** Survival curves (b), lymphocyte counts (c), and RABV neutralizing antibody titers (d) of ferrets infected with recCDV-5804p (Fig. 1), recCDV-5804p NΔ425-479, or recCDV-5804p Nectin 4-blind. **e,** Schematic of therapeutic efficacy studies utilizing recCDV-5804p NΔ425-479. **f-h,** Survival curves (f), CDV nasal lavage titers (g), and lymphocyte counts (h) of ferrets from (e). **i-j,** RABV (i) and pdmCA09 (j) nAb titers from ferrets shown in (e), determined using a single-cycle RABV reporter virus or pdmCA09, respectively. **k,** pdmCA09 nasal lavage titers of ferrets from (e) after challenge with pdmCA09 following recovery from CDV disease. Symbols (g-k) represent geometric means ± geometric SD, lines intersect means; log-rank (Mantel-Cox) test, median survival is stated (b, f); 2-way ANOVA with Sidak’s (c) or Dunnett’s (k) post-hoc test; mixed-effect analysis with Dunnett’s post-hoc test (g-j); n numbers as specified.

Clinical signs caused by the different recombinant strains, which were all in an otherwise identical recCDV-5804p genetic background, varied from severe for recCDV NΔ425-479, similar to the parental recCDV, to unremarkable for recCDV Nectin-4-blind (Extended Data Fig. 3a,b; Supplementary Fig. S4a-d). All recCDV Nectin-4-blind and a majority of recCDV NΔ425-479-infected animals survived, whereas ferrets inoculated with unmodified recCDV died within 12 days of infection (Fig. 2b). Surviving animals mounted a robust α-CDV nAb response within three weeks of infection (Supplementary Fig. S4e). PBMC-associated primary viremia titers of all three strains were statistically identical in the first ∼2 weeks after infection until animals inoculated with parental recCDV had succumbed (Extended Data Fig. 3c). Subsequently, viremia in recCDV Nectin-4-blind animals fully resolved, whereas ferrets infected with recCDV NΔ425-479 entered a 3-week low-level secondary viremia phase. Only animals inoculated with recCDV NΔ425-479 experienced lymphocytopenia with initial kinetics similar to that of the recCDV-infected group (Fig. 2c), whereas no statistically significant changes in lymphocyte counts were detected in ferrets infected with recCDV Nectin-4-blind, confirming over-attenuation of this recombinant. In the recCDV NΔ425-479-infected animals, lymphocytopenia fully resolved 4 weeks after infection. Pre-existing humoral α-RABV immunity collapsed within 3 weeks of recCDV NΔ425-479 infection and was not restored over a 3-month post-recovery period (Fig. 2d), whereas recCDV Nectin-4-blind caused only a temporary moderate decline in α-RABV nAbs titers that spontaneously resolved within 4 weeks of infection.

A comparison of clinical signs after infection of ferrets with the three recCDV strains with the presentation of human measles revealed that recCDV Nι1425-479 disease recapitulates hallmarks of human morbillivirus disease, albeit with accelerated onset of clinical signs (Extended Data Fig. 3d). Thus, we have developed a relevant surrogate animal model of human measles that strongly supports the morbillivirus immune amnesia hypothesis.

### Treatment at the onset of clinical signs alleviates morbillivirus immune amnesia

To explore the degree to which primary clinical signs of morbillivirus disease and immune amnesia can be mitigated pharmacologically, we initiated oral treatment with GHP-88309 at four discrete disease stages after infection with recCDV Nι1425-479: first onset of fever (5 dpi), rash (7 dpi), severe disease with diarrhea and conjunctivitis (10 dpi), and the first fatality in the vehicle group (24 dpi) (Fig. 2e). Treatment was continued b.i.d. until α-CDV nAbs became detectable in serum samples. In addition to following α-RABV immunity, we administered quadrivalent influenza vaccine to all animals 28 days prior to infection with CDV (–28-day prime, –14-day boost). Animals were monitored for 5 months post-CDV and the response to broad innate immune stimulation through intramuscular (i.m.) flagellin, infection with influenza virus A/CA/07/2009 (H1N1) (pdmCA09) and A/WI/67/2005 (H3N2), and homotypic rechallenge with pathogenic recCDV-5804p determined 8, 9.5, 12.5, and 18.5 weeks, respectively, after the original CDV infection.

All GHP-88309-treated animals and all ferrets of a CDV-vaccinated reference group survived until study end without developing any clinical signs, whereas approximately 40% of ferrets in the vehicle group succumbed to recCDV Nι1425-479 within 26 days of infection (Fig. 2f; Extended Data Fig. 4a,b). Oral GHP-88309 started 5 dpi significantly reduced CDV viremia titers (Extended Data Fig. 4c), but later initiation of treatment did not affect severity or duration of primary viremia. However, first administration of GHP-88309 up to 10 dpi statistically significantly shortened virus shedding into nasal lavages (Fig. 2g). Treatment started at the end of acute disease (24 dpi) did not modulate height or duration of viremia or virus shedding. Lymphocytopenia was suppressed in ferrets of the 5 dpi GHP-88309 group, resembling CDV-vaccine-mediated protection, and alleviated in the 7 dpi GHP-88309 group (Fig. 2h; Supplementary Fig. S5a-d). In animals first treated later than 7 dpi, lymphocytopenia was severe, closely resembling that in the vehicle group, and lymphocyte counts regained pre-infection levels only ∼10 weeks after CDV infection. Studies in nonhuman primates infected with MeV have revealed that viral RNA remains detectable in PBMCs up to 90 days after infection^16^.

Recapitulating this phenotype, we detected CDV RNA for over 64 days in PBMCs extracted from ferrets of all groups, except for CDV-vaccinated animals and ferrets first treated with GHP-88309 5 or 7 dpi (Extended Data Fig. 4d). Early treatment onset thus accelerated viral clearance and prevented persistence of viral RNA, which correlated with statistically significantly earlier appearance of α-CDV nAbs (Extended Data Fig. 4e). Whole genome sequencing of viruses recovered from GHP-88309-experienced animals 8-37 dpi with CDV did not reveal emergence of GHP-88309-characteristic^6^ resistance mutations (Supplementary Dataset S1).

Upon stimulation of animals with 50 µg flagellin i.m. post-recovery from acute CDV, we monitored expression of a panel of pro-inflammatory markers by PBMCs harvested before, or 1, 12, and 24 hours after, stimulation (Supplementary Fig. S6). Body temperature was increased in all animals after flagellin injection without significant differences in marker expression between groups (Extended Data Fig. 4f), indicating that innate response to an immunogen was not compromised after recovery from acute CDV disease.

Animals of all groups had robust α-RABV (Fig. 2i) and α-IAV (Fig. 2j) humoral immunity prior to CDV infection. Treatment with GHP-88309 5 dpi fully preserved nAbs titers, identical to the protective effect of CDV vaccination. Animals in all later treatment groups experienced a rapid decline in α-RABV and α-IAV nAbs titers over a 30-40 day period post-CDV infection, which was indistinguishable from that seen in the vehicle group and, in the case of α-RABV immunity, irreversible. However, low levels of α-IAV nAbs were present in the GHP-88309 7– and 10-dpi treatment groups at the time of challenge with pdmCA09 9.5 weeks after CDV infection. Accordingly, animals in both the 5– and 7– dpi GHP-88309 groups showed significantly reduced peak shed influenza virus load (Fig. 2k) and alleviated clinical signs of pdmCA09 infection (Extended Data Fig. 5a,b). All animals infected with pdmCA09 mounted a robust neutralizing response 2 weeks after challenge, indicating that CDV infection erased pre-existing immunity, but had no lasting suppressive effect on humoral immune competence to a new challenge post-recovery.

Since α-pdmCA09 nAbs are not cross-protective against subtype H3N2 IAVs, they can only provide protection by heterosubtypic immunity^17^. We therefore inoculated animals with an A/WI/67/2005 (H3N2) 3 weeks after the pdmCA09 challenge (87 dpi with CDV) to assess cell-mediated immune competence. As expected, ferrets of the CDV vaccine and 5-dpi GHP-88309 treatment groups had retained high α-H3N2 nAbs titers induced by the quadrivalent influenza vaccine (Extended Data Fig. 5c). Humoral α-H3N2 immunity in all other groups collapsed. None of the groups showed significant clinical signs (Extended Data Fig. 5d,e), α-H3N2 nAbs were rapidly rebuilt within 28 days of A/WI/67/2005 (H3N2) challenge (Extended Data Fig. 5f), and only low levels of virus shedding were detectable 24 hours after infection (Extended Data Fig. 5g), suggesting equivalent cross-reactive cell-mediated immunity derived from the pdmCA09 challenge in animals of all groups, which is consistent with previous reports for both the ferret model and human patients^18–22^. All animals furthermore mounted a robust humoral α-CDV response post-recovery and were fully protected against homotypic challenge with non-attenuated recCDV-5804p 130 days after the original CDV infection (Supplementary Fig. S7).

These results indicate that treatment initiated at the onset of first clinical signs of morbillivirus disease (fever; 5 dpi in the CDV ferret model) fully protects, while treatment started at the onset of rash (7 dpi) partially preserves, pre-existing immunity. Later onset of treatment improves outcome of morbillivirus disease but does not subvert immune amnesia.

### Fatal lung disease after unrelated respiratory virus infection followed by morbillivirus invasion

Naturally acquired IAV immunity is more robust than vaccine-induced protection^23^. To better mimic such immunity, we developed a consecutive infection ferret model that establishes a prior disease history in the animals (Fig. 3a). Influenza virus-naïve ferrets were inoculated 28 days before CDV infection with pdmCA09, with the intent to be followed for 5 months including challenge with pdmCA09 and CDV. Animals in reference groups were CDV vaccinated, or not infected with pdmCA09, respectively. Ferrets in all influenza virus groups developed fulminant clinical signs of IAV infection (Supplementary Fig. S8) and reached peak shed viral loads of 10^5^-10^6^ TCID_50_ units/ml nasal lavage 2 dpi (Extended Data Fig. 6a). pdmCA09 shedding ceased and all clinical signs resolved by 6 dpi, and ferrets had mounted robust humoral α-IAV (H1N1) immunity (Supplementary Fig. S9) when infected with attenuated recCDV NΔ425-479 4 weeks later (study day 0).

**Figure 3.**
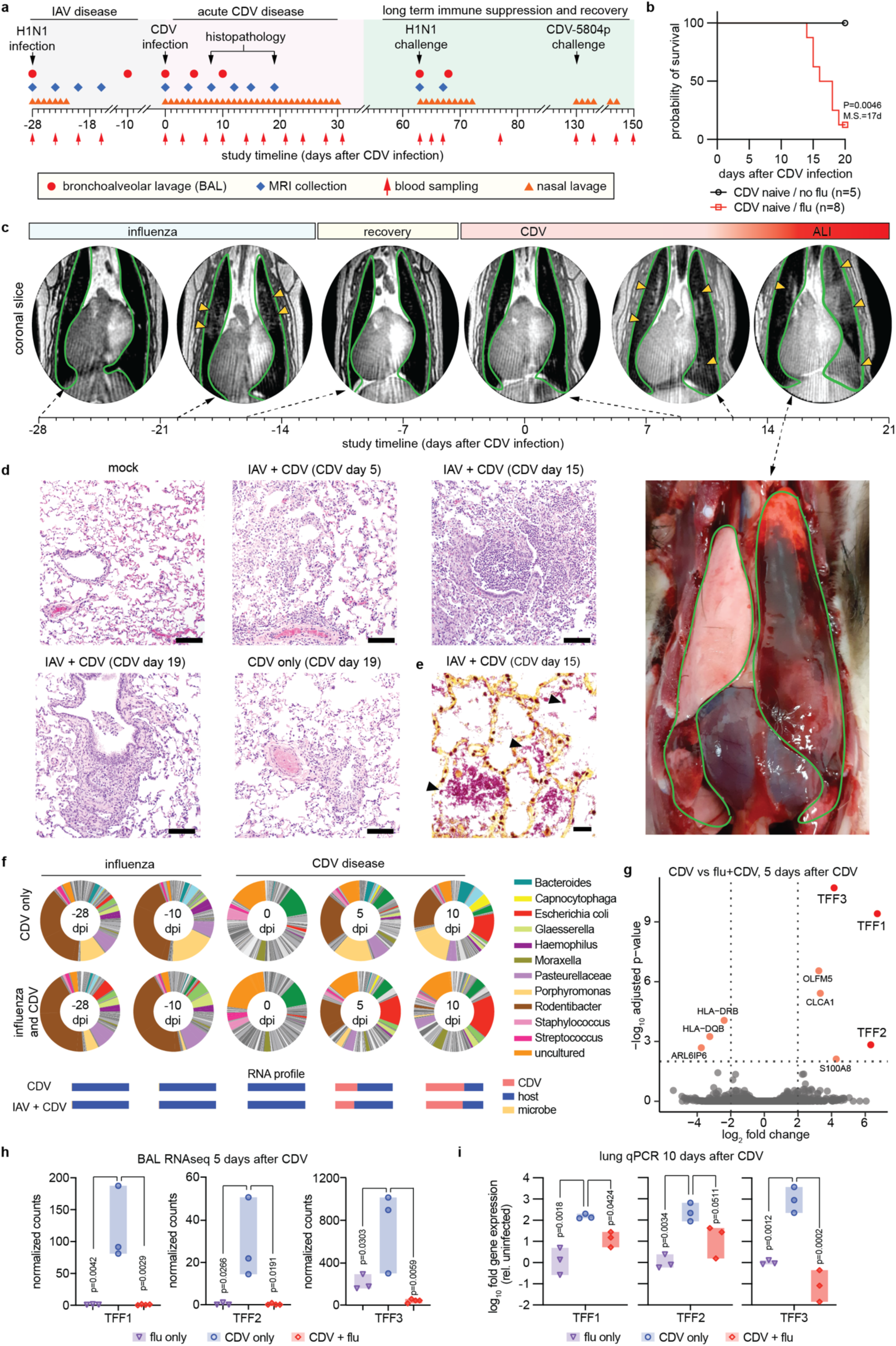
Fatal lung disease in ferrets infected with respiratory viruses prior to CDV infection. a, Study schematic. **b,** Survival curves of ferrets from (a). log-rank (Mantel-Cox) test, median survival is stated; n numbers as specified. **c,** MRI timeline performed on ferrets during primary pdmCA09 (0, 4, 8, and 12 dpi with influenza) and subsequent recCDV-5804p NΔ425-479 infection (8, 12, and 15 dpi with CDV). Lungs are outlined in green (dark appearance in MRI images), arrowheads denote fluid accumulation. Bottom right: necropsy of the ferret MRI-imaged on D15. **d,** Histopathology of ferret lungs from (a), extracted 5, 15, or 19 dpi with CDV. **e,** Gram staining of lung samples taken 15 dpi with CDV showing Gram-negative bacterial pneumonia. **f,** Metagenomics analysis of bacterial transcripts in BAL fluids collected from consecutively infected or CDV-only ferrets. Below each pie graph, RNA profiles of the relative composition of transcripts are shown. **g,** RNAseq screen of differentially expressed transcripts present in BAL fluids extracted 5 dpi with CDV from consecutively infected versus CDV-only ferrets. Significance thresholds (adjusted p<0.01, log2 fold change > 2) are shown as dotted horizontal and vertical lines, respectively. **h,** Trefoil factor (TFF1, TFF2, and TFF3) transcripts in BAL fluids harvested 5 dpi with CDV from consecutively infected versus CDV-only ferrets. **i,** RT-qPCR quantitation of relative presence of trefoil factor-encoding message in lung tissue of consecutively infected versus CDV-only ferrets, extracted 10 dpi with CDV. Symbols (h, i) represent individual animals, bars show range; 1-way ANOVA with Tukey’s (h) or Dunnett’s (i) post-hoc test; n=3.

Unexpectedly, most animals consecutively infected with this non-lethal IAV and low-lethal recCDV succumbed to acute hemorrhagic pneumonia with extensive spontaneous bleeding 14-18 dpi (Fig. 3b). Fatalities did not coincide with enhanced PBMC-associated CDV viremia titers or prolonged CDV shedding (Extended Data Fig. 6b,c). Postmortems of moribund vehicle-treated animals that presented with severe respiratory distress revealed major lung tissue injury with inflammatory lesions and wide-spread edema affecting several lobes (Extended Data Fig. 6d).

We employed pulmonary ferret MRI for longitudinal non-invasive assessment of lung disease progression (Supplementary Movie S1,S2). Transient viral pneumonia with localized fluid accumulation was present 8 days after pdmCA09, but fully resolved by 12 dpi (Fig. 3c). Infection of ferrets without prior influenza history with recCDV NΔ425-479 did not cause pulmonary edema. In contrast, pdmCA09-experienced animals developed pneumonia that was first detectable 10 days after CDV and rapidly advanced to major hemorrhage 14-19 dpi (Fig. 3c; Supplementary Movie S3, S4; Supplementary Fig. S10).

Histopathology of lung tissue extracted in repeat studies when vehicle-treated pdmCA09 and recCDV NΔ425-479-infected animals became moribund (15 and 18 dpi, respectively) revealed widespread edema, necrosis, and vasculitis in the small airways (Fig. 3d; Supplementary Fig. S11). Lung samples extracted 19 dpi from ferrets that were infected only with recCDV NΔ425-479 contained areas of inflammatory cellular infiltrates, bronchial epithelial hyperplasia and epithelial sloughing consistent with mild interstitial viral pneumonia, but showed no signs of edema and vasculitis, reflected by significantly lower pathology scores than those of the consecutively-infected group (Extended Data Fig. 7a). Lung tissues of CDV-vaccinated consecutively infected animals were unremarkable, their pathology scores resembling those of mock-infected ferrets. Gram-staining of tissue samples from consecutively-infected animals showed fulminant bacterial superinfections that were absent from animals infected with CDV only (Fig. 3e; Supplementary Fig. 12), which clinical microbiology identified as part of the commensal microbiome (Extended Data Fig. 7b). Metagenomics furthermore revealed a relative expansion of the *Escherichia* population in the lung microbiome 10 dpi with CDV in both singly and consecutively infected animals (Fig. 3f; Supplementary Dataset S2).

To assess whether priming for exacerbated disease is correlated with severity of the initial viral pneumonia, is strictly dependent on the 28-day interval between infections, or is IAV-specific, we treated the initial pdmCA09-infected animals with the broad-spectrum nucleoside-analog antiviral 4’-FlU (EIDD-2749)^24^, infected animals with pdmCA09 67 days before CDV, or inoculated ferrets with RSV-A2-L19F^25^ instead of pdmCA09, respectively (Extended Data Fig. 8a).

Bronchoalveolar lavages (BALs) were sampled at 5 predefined time points throughout the study. Consistent with previously demonstrated α-IAV efficacy of 4’-FlU^26^, once daily (q.d.) oral administration at 2 mg/kg bodyweight initiated 24 hours after infection with pdmCA09 statistically significantly shortened viral shedding and alleviated clinical signs (Extended Data Fig. 8b,c). However, pharmacological mitigation of IAV disease did not significantly alter viremia titers or virus shedding after subsequent infection with recCDV NΔ425-479 (Extended Data Fig. 8d,e) and the majority of animals succumbed to hemorrhagic pneumonia (Extended Data Fig. 8f). An expanded 67-day interval between IAV and CDV infections had no major alleviating effect on CDV disease presentation and hemorrhagic pneumonia outcome (Extended Data Fig. 8g-i).

Inoculation of ferrets with RSV-A2-L19F resulted in productive infection, characterized by efficient RSV shedding into nasal lavages (Extended Data Fig. 9a), but animals did not display overt clinical signs (Extended Data Fig. 9b). No cross-protective immunity with paramyxovirus infection was observed when we inoculated ferrets 28-days after RSV with recCDV NΔ425-479 (Extended Data Fig. 9c-e), and the majority of animals again succumbed to hemorrhagic pneumonia (Extended Data Fig. 9f).

### Differential expression of TFF peptides after CDV infection as a consequence of prior disease history

RNA-qPCR analysis of BAL samples taken during and after IAV infection of ferrets for expression levels of selected pro-inflammatory and wound-healing cytokines relative to those prior to exposure revealed that respiratory tissues of IAV-experienced animals were in an anti-inflammatory state at the time of CDV infection compared to immune homeostasis of IAV-naïve animals (Supplementary Fig. S13a-e).

Comparative RNAseq screening of BAL samples extracted 5 dpi from ferrets infected with recCDV NΔ425-479, or consecutively infected with pdmCA09 followed by recCDV NΔ425-479, identified 9 genes as significantly differentially expressed between study groups (Fig. 3g). These included all three genes encoding trefoil factor (TFF) peptides (Fig. 3h), which are known to be involved in protection and repair of gastrointestinal^27^ and, although less well studied, respiratory^28^ epithelia. Follow-up RT-qPCR of lung tissue extracts harvested 10 days after CDV infection confirmed that TFF-encoding mRNAs were upregulated several 100-fold (TFF1 and 2) to 1,000-fold (TFF3) in animals infected only with CDV compared to the consecutively infected groups (Fig. 3i). pdmCA09 infection alone resulted in only slightly increased TFF1 and 2 expression 10 dpi and TFF3 was not upregulated (Extended Data Fig. 10a-c), indicating that extreme induction of TFF1 and 3 is morbillivirus-specific rather than a general response to respiratory virus infections. In the mouse respiratory tract, TFF1 and 3 reportedly colocalize with mucins Muc5AC and Muc5B^29^. However, RT-qPCR did not reveal an equivalent differential upregulation of either Muc5 expression level in the different infection groups (Supplementary Fig. S14a,b).

These results demonstrated that immune priming through unrelated primary viral pneumonia sets the stage for exacerbated subsequent morbillivirus disease. CDV infection during, or shortly after, the recovery phase of an inflammatory episode alters the quality of the host defense to morbillivirus disease, resulting in significantly lower expression of respiratory epithelium protective TFFs, which emerged as correlative for risk to advance to hemorrhagic pneumonia.

### Late onset treatment of CDV infection does not mitigate signs of CDV disease, but alters lethal outcome

To assess the effect of treatment of morbillivirus infection on disease outcome, we again consecutively infected animals with pdmCA09 followed by recCDV NΔ425-479 (Supplementary Fig. S15), and initiated oral treatment with GHP-88309 5, 7, 10, and 14 dpi with CDV (Fig. 4a). Consistent with results of the previous GHP-88309 treatment studies, PBMC-associated CDV viremia was mitigated only when treatment was started 5 dpi (Fig. 4b). All animals started on GHP-88309 5 or 7 dpi with CDV survived, whereas vehicle-treated animals succumbed to hemorrhagic pneumonia between days 14-19 after CDV infection (Fig. 4c). A majority of animals in the 10 dpi treatment group recovered, but most ferrets first treated with GHP-88309 14 days after CDV developed lethal pneumonia. Lymphocyte counts were fully preserved in CDV-vaccinated control animals and ferrets of the 5 dpi treatment group, and were partially preserved in animals of the 7 dpi group (Extended Data Fig. 10d). Although later treatment start did not mitigate lymphocytopenia, PBMC repopulation was accelerated in the 10 dpi treatment group. Circulating granulocyte populations did not deviate from the normal range in any study group, making granulocytes transiently the predominant white blood cell population in animals experiencing lymphocytopenia (Supplementary Fig. S16). nAb titers against RABV and IAV H1N1 reflected the lymphocytopenia results; fully preserved in vaccinated and 5 dpi GHP-88309-treated animals, partially preserved in the 7 dpi treatment group, and not preserved when GHP-88309 was initiated 10 or 14 dpi with CDV (Fig. 4d,e). Accordingly, only ferrets in the 5 and 7 dpi GHP-88309 treatment groups were fully or partially, respectively, protected against pdmCA09 challenge 62 days after CDV (Fig. 4f). However, surviving animals of all GHP-88309 treatment groups mounted a robust *de novo* α-IAV H1N1 nAb response following challenge with pdmCA09 (Extended Data Fig. 10e). At study end, all ferrets had furthermore developed robust immunity against CDV reinfection (Supplementary Fig. S17).

**Figure 4.**
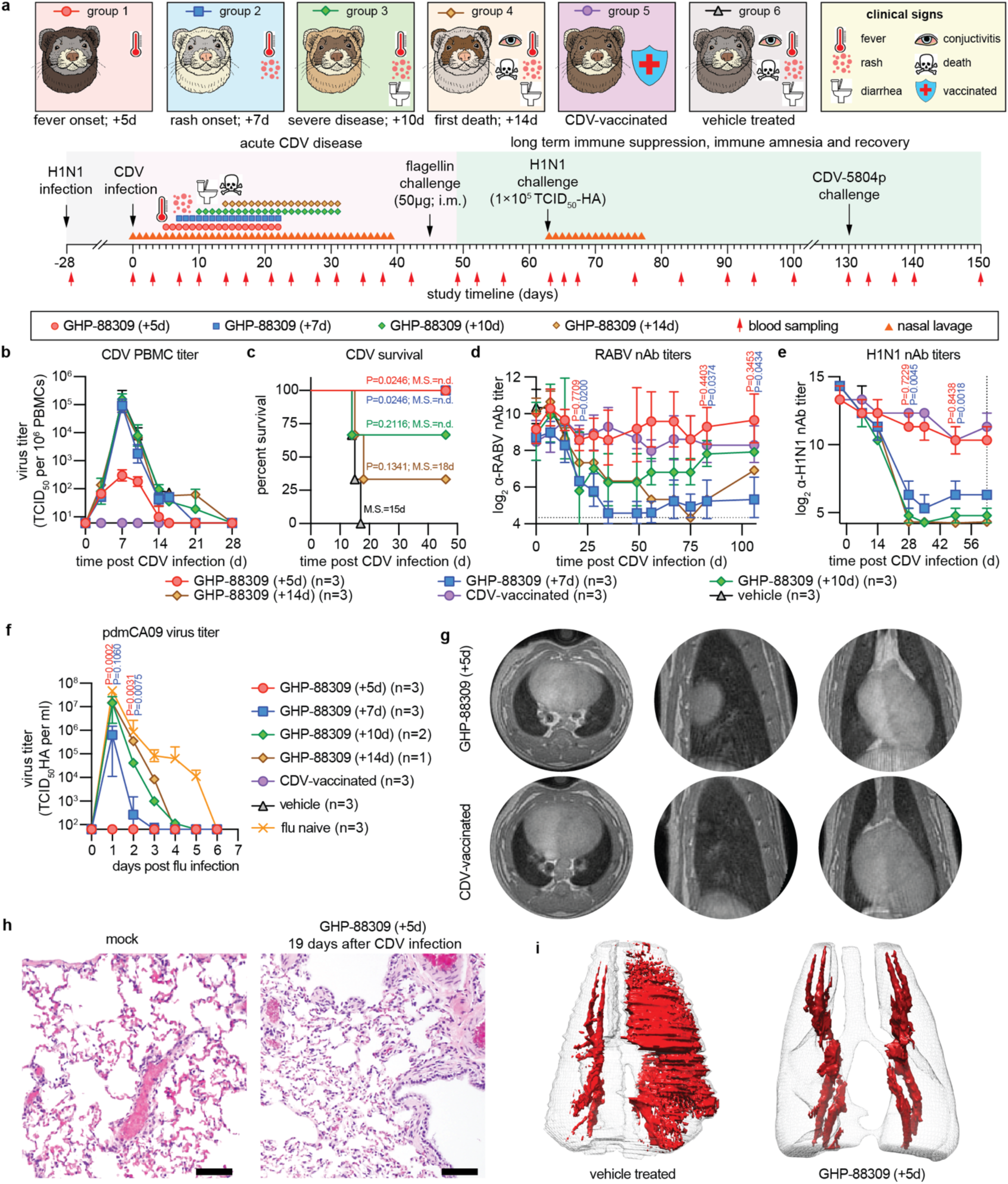
Effect of therapeutic intervention on severe disease outcomes. a, Study schematic. **b-c,** PBMC-associated viremia titers (b) and survival (c) of consecutively infected and GHP-88309-treated ferrets from (a). **d-e,** RABV (d) and IAV H1N1 (e) nAb titers mounted by animals from (a). **f,** pdmCA09 nasal lavage titers after IAV challenge of ferrets recovered from CDV disease. **g,** MRI slices of consecutively infected and GHP-88309-treated (top rww) or CDV vaccinated (bottom row) ferrets, taken 15 dpi with CDV. Axial, coronal, and sagittal slices of the same lung are shown. **h,** Lung histopathology of consecutively infected and GHP-88309-treated or uninfected (mock) ferrets, assessed 19 dpi with CDV. **i,** 3D MRI reconstructions and segmentations of lungs from consecutively IAV CDV-infected and GHP-88309 or vehicle-treated ferrets. Images were collected 15 dpi with CDV; fluids shown in red. Symbols (b, d, e, f) represent geometric means ± geometric SD, lines intersect means; log-rank (Mantel-Cox) test, median survival is stated (c); 2-way ANOVA with Dunnett’s post-hoc test (d-f); n numbers as specified.

Longitudinal MRI analysis 15 dpi with CDV of consecutively IAV and CDV-infected animals treated with GHP-88309 demonstrated suppression of pulmonary edema and hemorrhage (Fig. 4g,h; Supplementary Movie S5,S6), which was confirmed by histopathology of treated animals (Fig. 4i). Again, whole genome sequencing of CDV recovered from GHP-88309-treated ferrets 9-19 dpi revealed no known^6^ resistance mutations (Supplementary Dataset S3).

Outcome reversal of lethal hemorrhagic pneumonia through late-onset GHP-88309 established precedent for therapeutic benefit of direct-acting antiviral therapy initiated after the time window for mitigation of primary clinical signs of morbillivirus disease has closed. The results demonstrate that GHP-88309 treatment paradigms for mitigation of morbillivirus-induced immune amnesia are independent of whether immunity was naturally acquired or vaccine-induced.

## Discussion

Reflecting a paucity of effective antiviral therapeutics, their benefit is poorly defined for most pathogenic paramyxoviruses and unknown for members of the *Morbillivirus* genus, specifically. Recent work testing remdesivir therapy in a non-human primate model of MeV infection reported transient reduction of viral RNA through post-exposure prophylactic treatment, but lymphocytopenia and clinical disease parameters were not improved and virus replication rebounded^30^. Since remdesivir was designed to yield high liver exposure^31^, sustained inhibitory concentrations may not be reached in MeV-relevant target cells and tissues. Having identified orally efficacious morbillivirus^13,32^ and broadened-spectrum paramyxovirus^6^ inhibitors, we established in this study treatment paradigms for morbillivirus disease. Results support five major conclusions: i) efficacious antivirals such as GHP-88309 expand the therapeutic time window to eight days compared to therapeutic vaccination; ii) infections with engineered attenuated recCDV confirm the morbillivirus immune amnesia hypothesis in the ferret model; iii) therapeutic mitigation of primary clinical signs of morbillivirus disease and immune amnesia is beneficial when treatment is initiated before, or at peak, of primary viremia; iv) prior disease history of the host defines the risk for exacerbated, fatal morbillivirus infection; and v) late-onset anti-morbillivirus treatment prevents lethal bacterial superinfection.

The impact of prior infection on the outcome of morbillivirus infection was most unexpected. Although heightened long-term susceptibility to secondary infections after severe primary viral or bacterial pneumonia was described in mice^33^, this sepsis-induced transient immunosuppression was attributed to poor antigen-presentation capacity of DCs and AMs in the TGF-Δ-driven anti-inflammatory microenvironment established during the tissue healing phase after resolution of the primary infection^34^. However, immune priming for catastrophic morbillivirus disease did not require severe primary pneumonia, since highly efficacious treatment of pdmCA09 infection with 4’-FlU^26^ did not prevent hemorrhagic pneumonia after subsequent CDV infection. Unlike therapeutic intervention with the primary IAV infection, late-onset treatment of the morbillivirus infection with GHP-88309 initiated shortly before uncontrolled amplification of commensal bacteria changed outcome, indicating that CDV replication in the respiratory epithelium after basolateral re-invasion of lungs is instrumental for progression to lethal secondary complications. Our study revealed that morbillivirus triggers massive upregulation of TFF1, TFF3 and, to a lesser degree, TFF2 expression, provided infection occurred during lung immune homeostasis. In the respiratory tract, TFFs are predominantly produced by glandular and mucus-secreting epithelial and hematopoietic cells^28,35–37^, and TFF1 and 3 specifically enhance mucociliary bacterial clearance through increasing mucus viscosity^28,38,39^.

Based on these observations, we propose an immune-priming-based mechanism leading to exacerbated disease. IAV infection reportedly results in transient population of the respiratory tract with functionally altered DCs and AMs to restore immune homeostasis after the pro-inflammatory response, which are unable to clear bacterial pathogens efficiently^34^. CDV infection during the recovery phase, followed by basolateral invasion of the respiratory epithelium after primary viremia, further depletes DCs and AMs^8^ and fails to upregulate TFF expression, setting the stage for catastrophic bacterial pneumonia through impaired clearance of commensal bacteria. How does prior IAV infection affect TFF1 and 3 upregulation? Type 2 cytokines IL-4 and IL-13 stimulate TFF3 synthesis in the GI tract in a STAT6-dependent process^28^, but regulation of TFF3 expression in the airway epithelium is poorly understood. In mice, IAV stimulates lung-resident lineage-negative epithelial progenitor (LNEP) cells that are the major responders in distal lung after tissue damage^40^ and express high levels of TFF2^28^ potentially creating a negative feedback loop that prevents TFF1 and 3 upregulation. Future work must focus on the regulation of TFF3 levels in the homeostatic respiratory tract to better understand the molecular basis for long-lasting impaired expression after influenza virus priming.

It is currently unknown whether MeV equally induces TFF1 and 3 expression in the human respiratory tract and what impact disease history of measles patients may have on severity of secondary complications. Anecdotal cases of hemorrhagic pneumonia associated with measles have been reported^41,42^, but prior disease history of the patients was not documented. However, bacterial superinfection after measles such as laryngitis, bronchitis, and otitis media are common^11^. Typically attributed to impaired adaptive immunity due to lymphocytopenia, our ferret data support that respiratory disease history of measles patients should be considered as risk factor for advance to severe bacterial superinfections. Limitations affecting the predictive power of the CDV ferret model for human measles include different disease dynamics^43^, higher mortality rates of the attenuated recCDV NΔ425-479, higher inherent neurotropism of CDV than MeV^43^, and untested cross-species consistency of pharmacokinetic and efficacy performance of GHP-88309.

This study establishes treatment paradigms for efficacious pharmacological intervention in morbillivirus disease, defines respiratory disease history as a correlate for the risk of severe bacterial superinfection, and provides precedent for therapeutic benefit of treatment of an acute RNA virus infection with direct-acting antivirals initiated after the window for mitigation of primary clinical signs has closed.

## Acknowledgements

We thank the Georgia State University Department of Animal Resources, the Georgia State University Advanced Translational Imaging Facility (ATIF), and the University of Georgia Pathogenesis Core for expert assistance. The VSV-ΔG and recCDV-5804p genomic plasmids and Vero-canine SLAM cells were kind gifts of M.J. Schnell, V. von Messling, and Y. Yanagi, respectively. This study was supported, in part, by public health service grants AI071002 (to R.K.P.) and AI171403 project 3 (to R.K.P.) and Scientific Core D (to A.L.G.) from the NIH/NIAID.

## Conflict of Interests

RKP and RMC are co-inventors on a patent filing covering method of use of GHP-88309 for antiviral therapy. This study could affect their personal financial status. RKP reports contract testing from Enanta Pharmaceuticals and Atea Pharmaceuticals, and research support from Gilead Sciences, outside of the described work. RMC reports consulting for Merck & Co, outside of the described work. ALG reports contract testing from Abbott, Cepheid, Novavax, Pfizer, Janssen and Hologic and research support from Gilead Sciences, outside of the described work. RdS reports research support from Gilead Sciences and Themis Biosciences, outside of the described work. All other authors declare that they have no competing interests to report.

## Methods

### Study design

The objectives of this study were to determine the efficacy of GHP-88309 against morbillivirus disease in the CDV-ferret animal model and identify therapeutic windows for the successful treatment of a lethal morbillivirus infection.

The CDV-ferret model was chosen because it provides a surrogate animal model of severe morbillivirus disease, including many of the hallmark symptoms and clinical signs of measles virus in humans. The effect of treatment on virus replication and clinical signs in ferrets was determined using multiple therapeutic treatment regimens. Treatment was considered efficacious when statistically significant reductions in viremia and shed virus titers in PBMCs and nasal lavages, respectively, were observed and the duration and severity of clinical signs and immune suppression were decreased. Study endpoints were predefined prior to initiating experiments. At least two ferrets were used in all PK and tolerability studies. Groups of at least three ferrets were used in all *in vivo* efficacy studies. Before initiating experiments, animals were randomly assigned into groups. Exact numbers of independent biological repeats (individual animals) for each experiment are specified in the figures or figure legends. All quantitative source data are provided in the Supplementary Dataset S4.

### Detailed experimental methods

Detailed methods describing cell lines and viruses used, transfections protocols, virus yield reduction assays, PK studies, animal infections, RNA extraction and RT-qPCR profiling, RNAseq screening, virus neutralization assays, and pulmonary ferret MRI are provided as supplementary information.

### Inclusion and ethics statement

All animal work was performed in compliance with the Guide for the Care and Use of Laboratory Animals of the National Institutes of Health and the Animal Welfare Act Code of Federal Regulations. Experiments involving ferrets were approved by the Georgia State University IACUC under protocols A22035 and A18035.

### Statistics and reproducibility

One-way or two-way analysis of variance (ANOVA) with Dunnett’s or Sidak’s multiple comparison post-hoc tests, was used to assess statistical differences. All statistical analyses were carried out in GraphPad Prism software (Version 9.3.1). Specific statistical tests applied to individual data sets are specified in the corresponding figure legends. The number of individual biological replicates for all graphical representations are shown in the figures. Representations of mean ±SD or median ± 95% CI of experimental uncertainty are shown and specified in the figure legends. Fourteen-day tolerability studies were based on two ferrets. All statistical analyses and exact P values are shown in the Supplementary Dataset S5. Alpha levels were set to 0.05 for all significance analyses.

### Reporting summary

Further information on research design is available in the Reporting Summary linked to this article.

## Data availability

The amplicon tiling sequencing reads generated in this study have been deposited in the NCBI BioProject database under accession code PRJNA1004336 (https://www.ncbi.nlm.nih.gov/bioproject/?term=PRJNA1004336). All other data generated in this study are provided in this article, the supplementary information, supplementary data files, and the Source Data file. Source data are provided with this paper.

## Code availability

Sequencing reads were analyzed using the TAYLOR pipeline (available at https://github.com/greninger-lab/covid_swift_pipeline). All commercial computer codes and algorithms used are specified in Methods.

## Extended Data Figures

**Extended Data Fig. 1.**
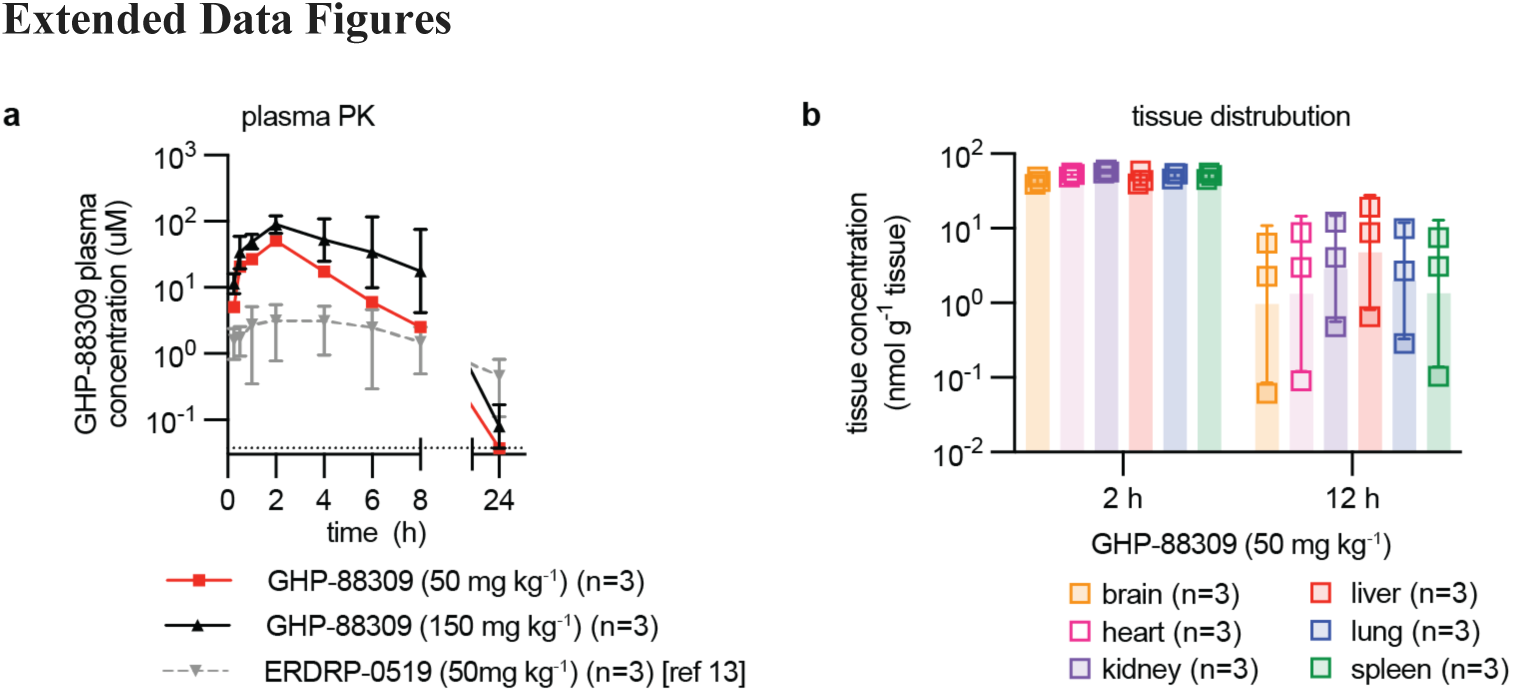
Single-dose oral PK study of GHP-88309 in ferrets. a, Plasma concentration of GHP-88309 determined over a 24-hour period after a single oral dose of 50 mg/kg (red squares) or 150 mg/kg (black triangles). For comparison, historical data^13^ of ERDRP-0519 plasma levels after a single oral dose of 50 mg/kg (grey triangles, dotted grey lines) are shown. **b,** GHP-88309 exposure in selected ferret organs at 2 and 12 hours after administration of GHP-88309 (50 mg/kg). Symbols represent geometric means ± geometric SD, lines intersect means (a), or independent biological repeats (b), columns denote geometric means ± geometric SD; n=3.

**Extended Data Fig. 2.**
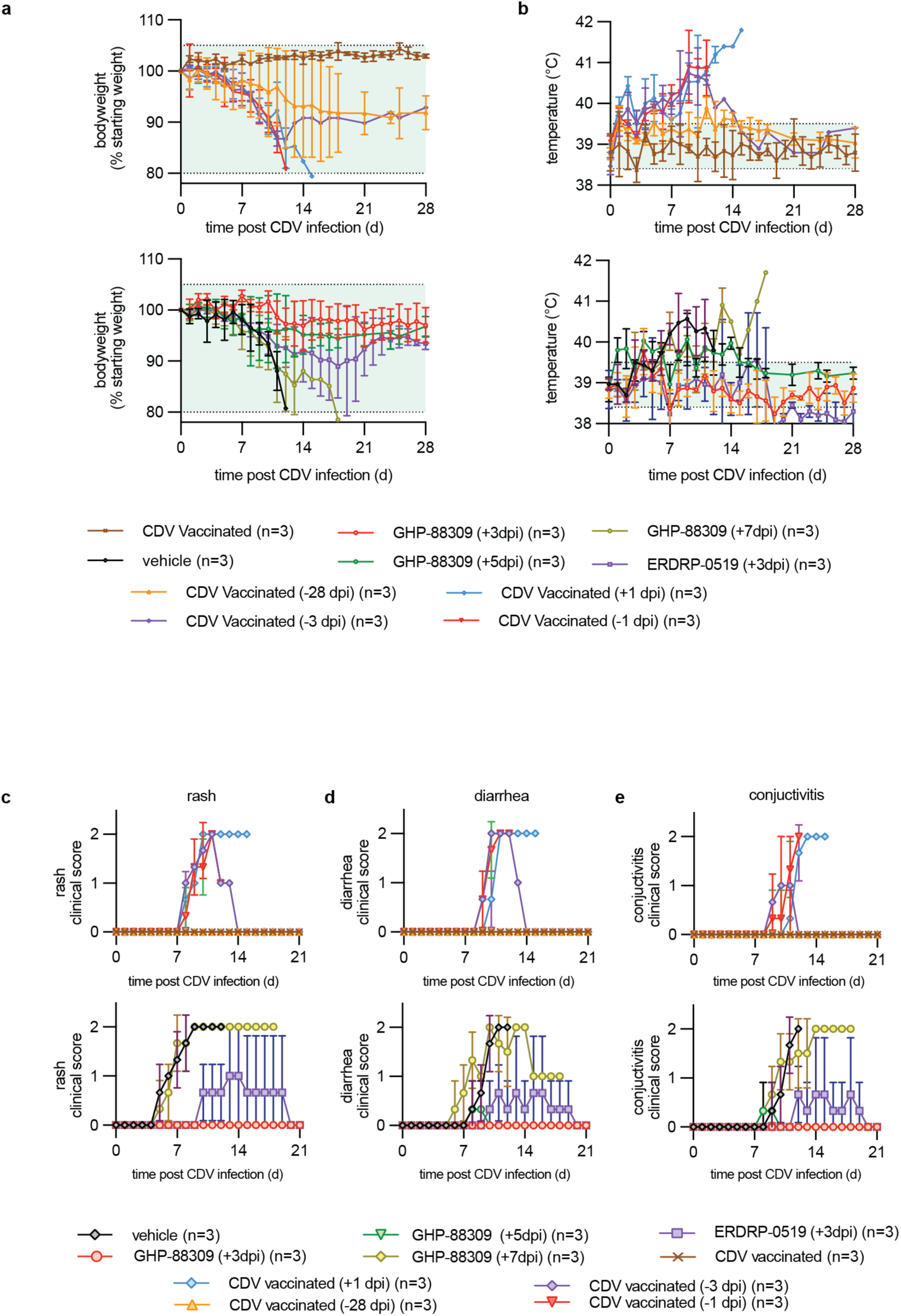
Clinical signs of ferrets infected with recCDV-5804p. a-e, Bodyweight (a), temperature (b), and clinical scores of rash (c), diarrhea (d), and conjunctivitis (e) of ferrets shown in Fig. 1. Symbols represent arithmetic means ± SD, lines intersect means; n=3.

**Extended Data Fig. 3.**
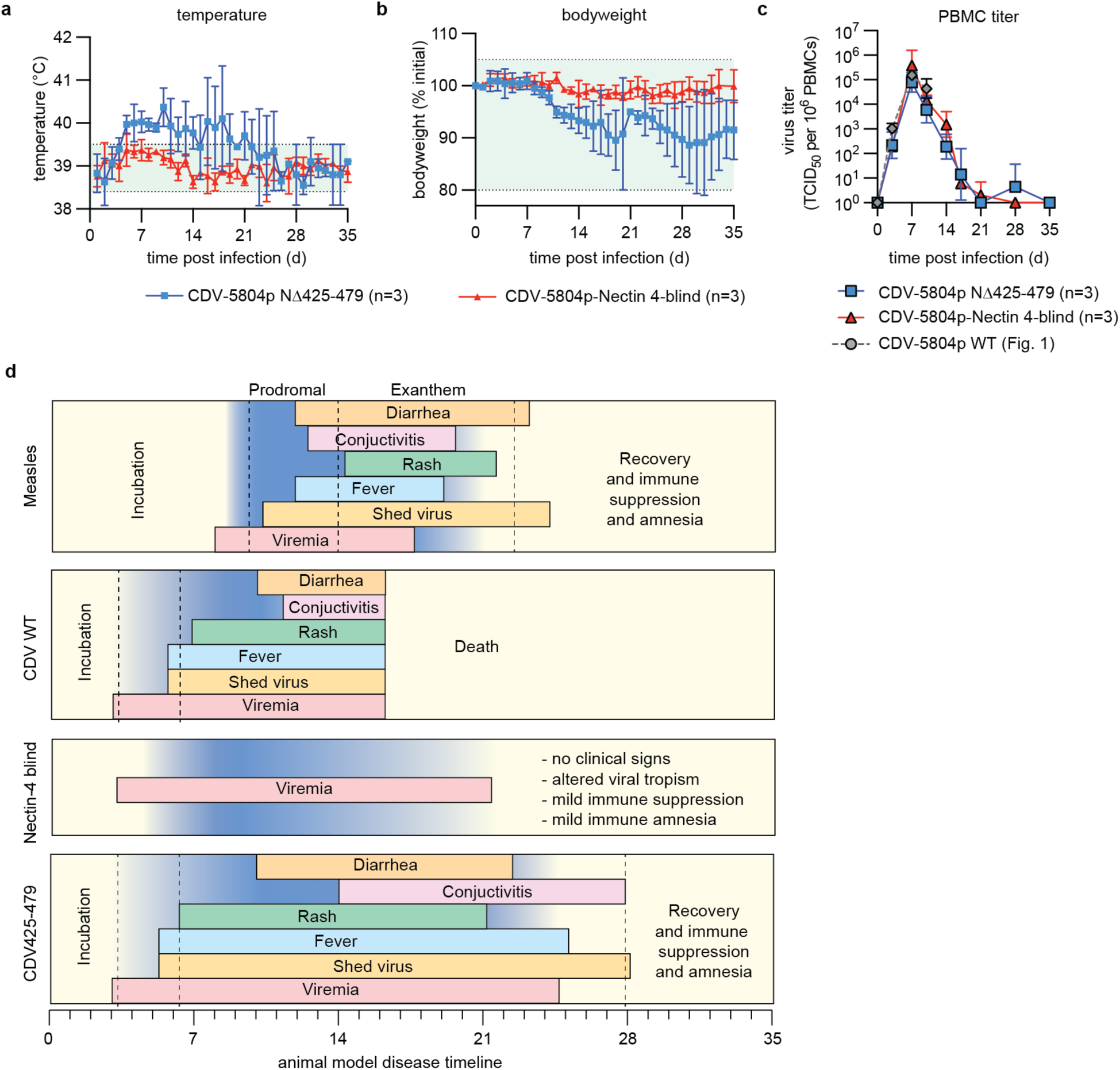
Clinical signs and viremia after infection with recCDV-5804p N/¢1425-479 versus recCDV-5804p-Nectin 4-blind. a-b, Temperature (a) and bodyweight (b). **c,** PBMC-associated viremia titers of ferrets, monitored for 35 days after infection of animals as specified in Fig. 2a. Symbols represent arithmetic (a, b) or geometric (c) means ± SD, lines intersect means; n=3. **d,** 2D schematic comparing disease dynamics and clinical signs of measles virus infections in humans, with wild-type recCDV-5804p, recCDV-5804p-Nectin 4-blind, and recCDV-5804p N/¢1425-479 infections in ferrets.

**Extended Data Fig. 4.**
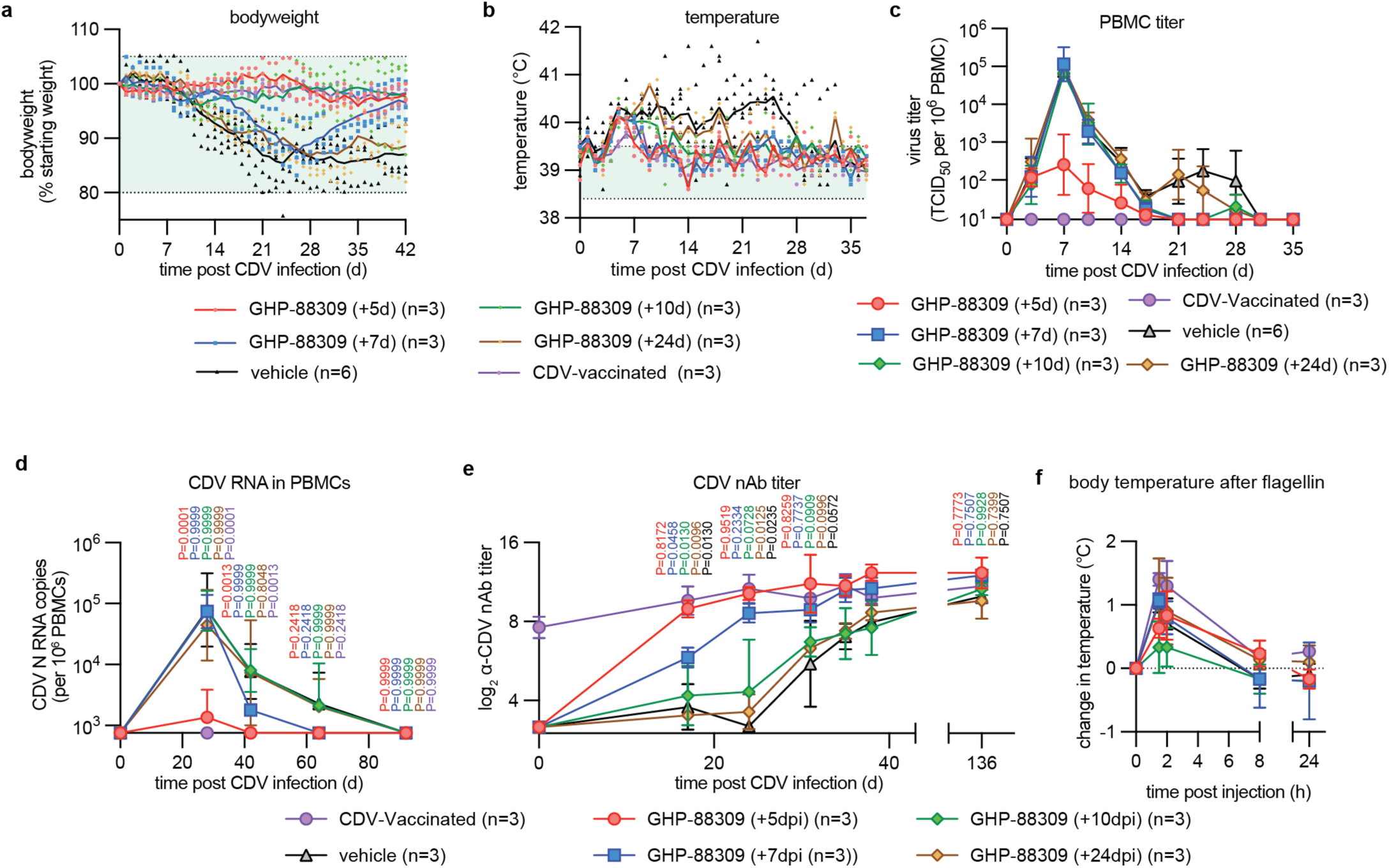
Clinical presentation and antiviral response of recCDV-5804p N/¢1425-479-infected and GHP-88309-treated ferrets. a-b, Bodyweight (a) and temperature measurements (b) assessed once daily over the course of acute CDV disease of ferrets from Fig. 2e. **c,** PBMC-associated viremia titers monitored until undetectable. **d,** Quantitation of CDV-specific N RNA in circulating PBMCs of ferrets monitored until undetectable. 2-way ANOVA with Dunnett’s post-hoc test. **e,** CDV nAb titers determined using CDV-5804p as target. Mixed-effects analysis with Dunnett’s post-hoc test. **f,** Changes in body temperature after i.m. administrations of flagellin as specified in Fig. 2e. Symbols represent individual animals, lines intersect arithmetic means (a, b), geometric means ± geometric SD c-e), or arithmetic means ± SD (f); green shading denotes normal range; n=3.

**Extended Data Fig. 5.**
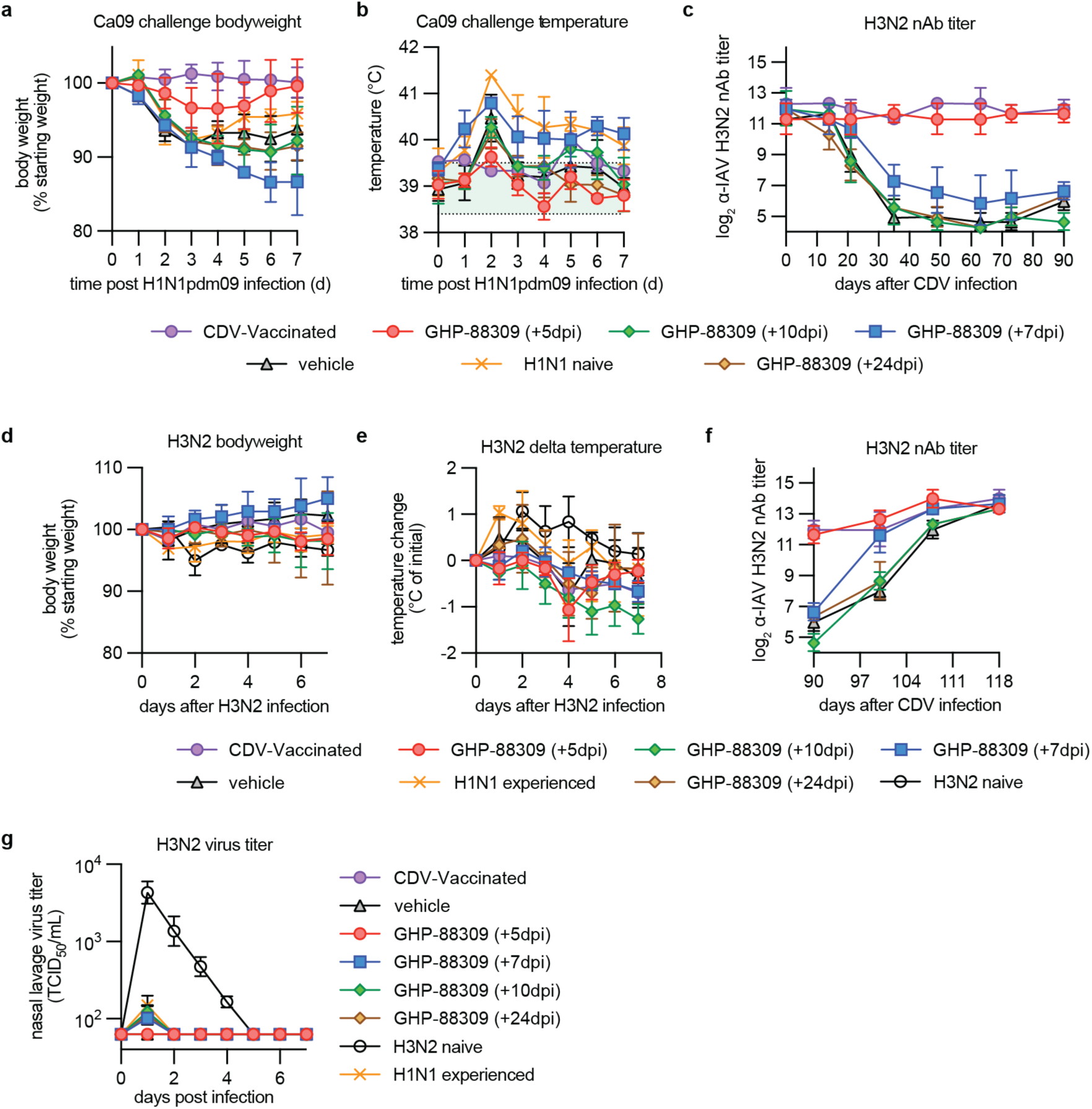
IAV challenge of ferrets after recovery from CDV disease. a-b, Bodyweight (a) and temperature (b) of ferrets after recovery from CDV disease and infection with pdmCA09 (H1N1) as specified in Fig. 2e; green shading denoted normal range. **c,** a-IAV H3N2 nAb titers in ferrets from Fig. 2e surviving CDV infection. **d-e,** Bodyweight and temperature of ferrets after infection with IAV-Wyo (H3N2) as specified in Fig. 2e. **f,** a-IAV H3N2 nAb titers infection with IAV-Wyo (H3N2). **g,** IAV-Wyo (H3N2) nasal lavage titers collected after challenge with IAV-Wyo. Symbols represent arithmetic means ± SD (a, b, d, e) or geometric means ± geometric SD (c, f, g); n=3.

**Extended Data Fig. 6.**
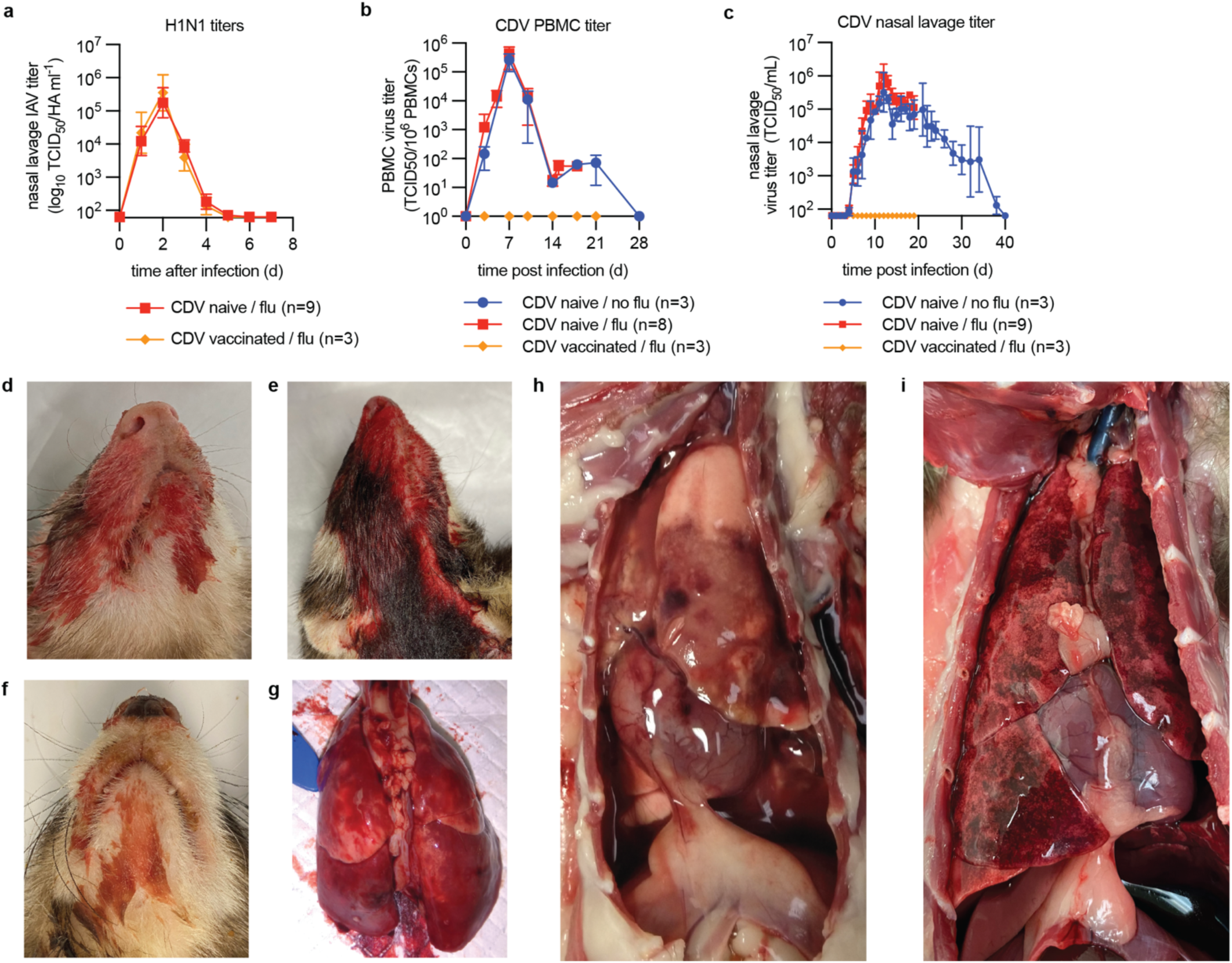
Exacerbated lung disease after consecutive infection of ferrets with IAV and CDV. a, pdmCA09 (H1N1) nasal lavage virus titers after primary IAV infection, 28 days prior to recCDV-5804p N/¢1425-479 as specified in Fig. 3a. **b, c**, PBMC-associated CDV viremia (b) and nasal lavage (c) titers after infection of ferrets with recCDV-5804p N/¢1425-479. Symbols in (a-c) represent geometric means ± geometric SD; n=3-9 as specified. **d-i,** Macroscopic (d-f) and necropsy (g-i) presentation of hemorrhagic pneumonia in consecutively infected ferrets. Enlarged views of gross pathology of ferrets shown in Fig 3c.

**Extended Data Fig. 7.**
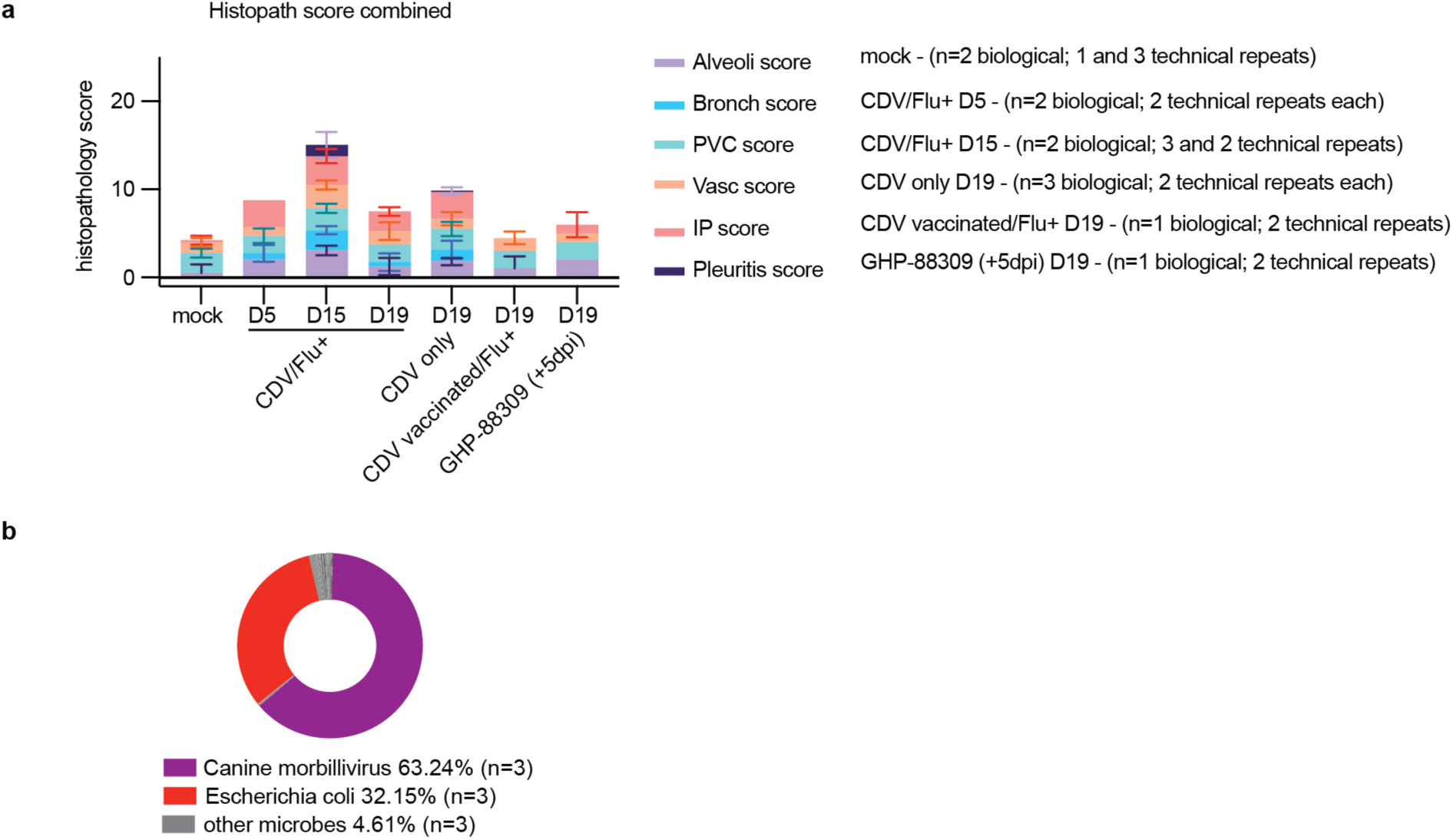
Histopathology scores and metagenomics after consecutive infection of ferrets. a, H&E-stained lung sections of ferrets from Fig. 3a extracted 5, 15, or 19 dpi with CDV were scored as specified in Supplementary Methods. Bars represent mean scores ± SD for each criterion, n numbers as specified; for clinical scoring, technical and biological repeats were considered equally for calculation of variance. **b,** Metagenomics analysis of lung tissues extracted from consecutively IAV and CDV-infected animals 15 (n=2) or 19 (n=1) dpi with CDV. Relative distribution of pathogen-specific reads is shown; n=3.

**Extended Data Fig. 8.**
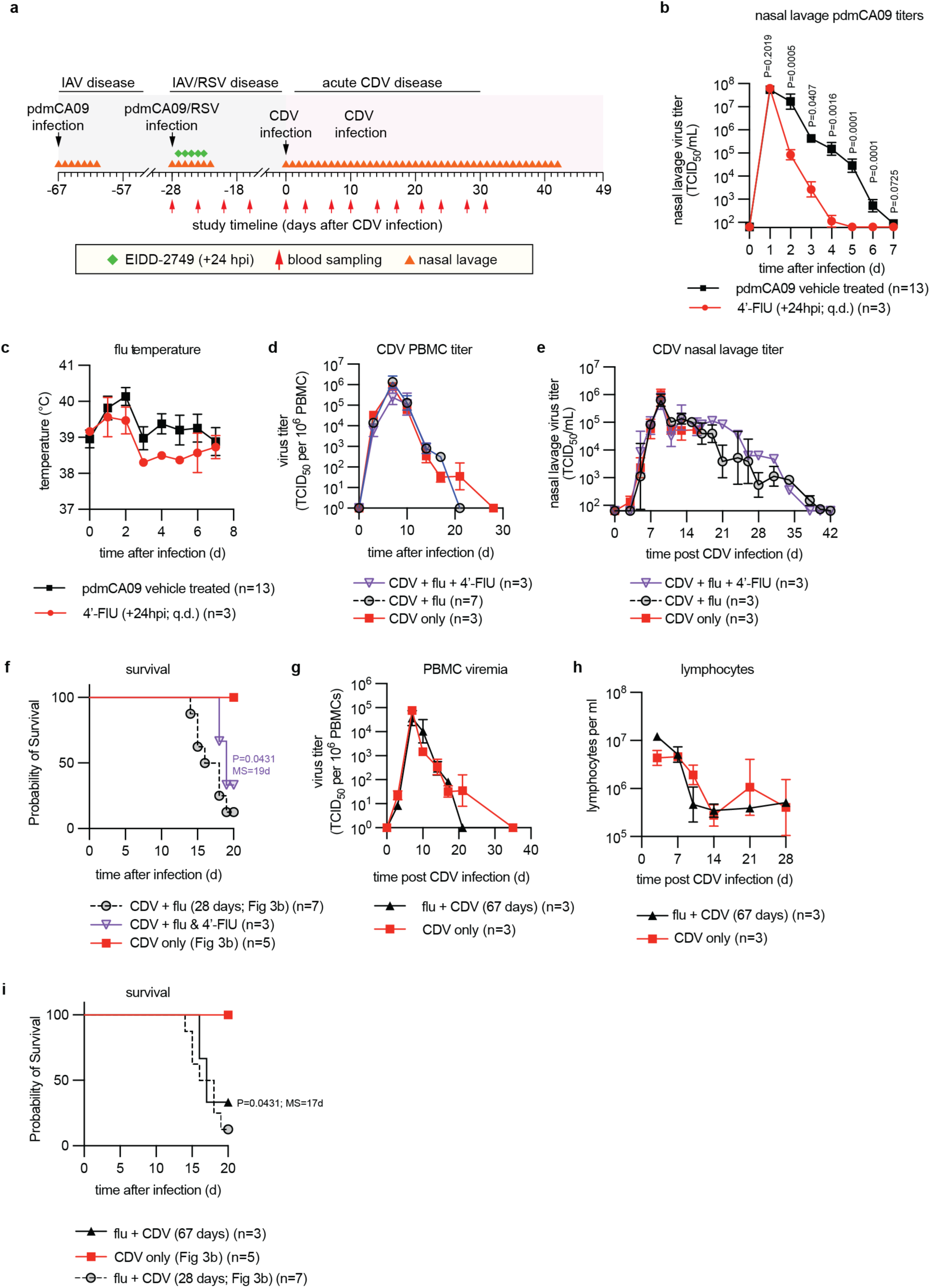
Effect of treating primary H1N1-pdmCA09 infection prior to CDV. a, **Schematic of study** design. **b-c,** pdmCA09 (H1N1) shed virus titers (b) and body temperature (c) of ferrets infected with pdmCA09 and treated with 4’-FlU or vehicle, 28 days prior to infection with recCDV-5804p N/¢1425-479 as shown in Fig. 3. **d-e,** PBMC-associated CDV viremia (d) and nasal lavage (e) titers of ferrets infected with recCDV-5804p N/¢1425-479. **f,** Survival of consecutively IAV CDV-infected ferrets after treatment of primary IAV with 4-FlU. **g-h,** PBMC-associated primary viremia titers (g), circulating lymphocyte counts (h), and survival (i) of ferrets infected with recCDV-5804p N/¢1425-479 67 dpi with primary pdmCA09. A control group of influenza-naïve ferrets was infected with recCDV-5804p N/¢1425-479 (red triangles). Symbols represent geometric means ± geometric SD (b, d, e, g, h) or arithmetic group means ± SD (c), lines intersect means; log-rank (Mantel-Cox) test, median survival is stated (f, i); n numbers as specified.

**Extended Data Fig. 9.**
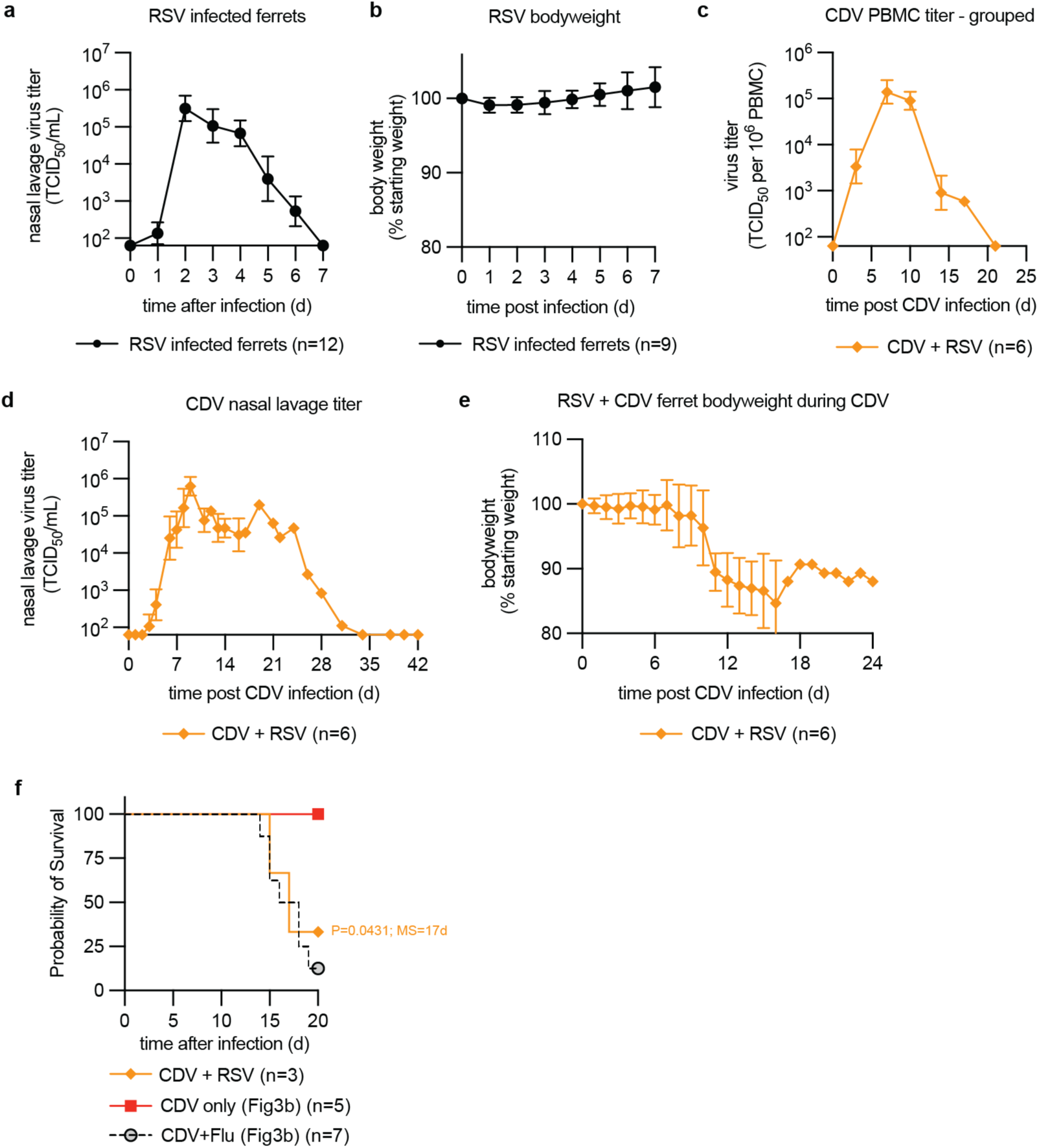
Exacerbated disease after primary infection with RSV followed by recCDV-5804p N/¢1425-479. a, RSV shed virus titers in nasal lavages. **b,** Bodyweight of ferrets after infection with RSV-A2-L19F. **c-f,** PBMC-associated CDV viremia (c) and nasal lavage (d) titers, animal bodyweight (e) and survival (f) after infection of ferrets with recCDV-5804p N/¢1425-479 28 dpi with RSV. Symbols represent geometric means ± geometric SD (a, c, d) or arithmetic means ± SD (b, e), lines intersect means; log-rank (Mantel-Cox) test, median survival is stated (f); n numbers as specified.

**Extended Data Fig. 10.**
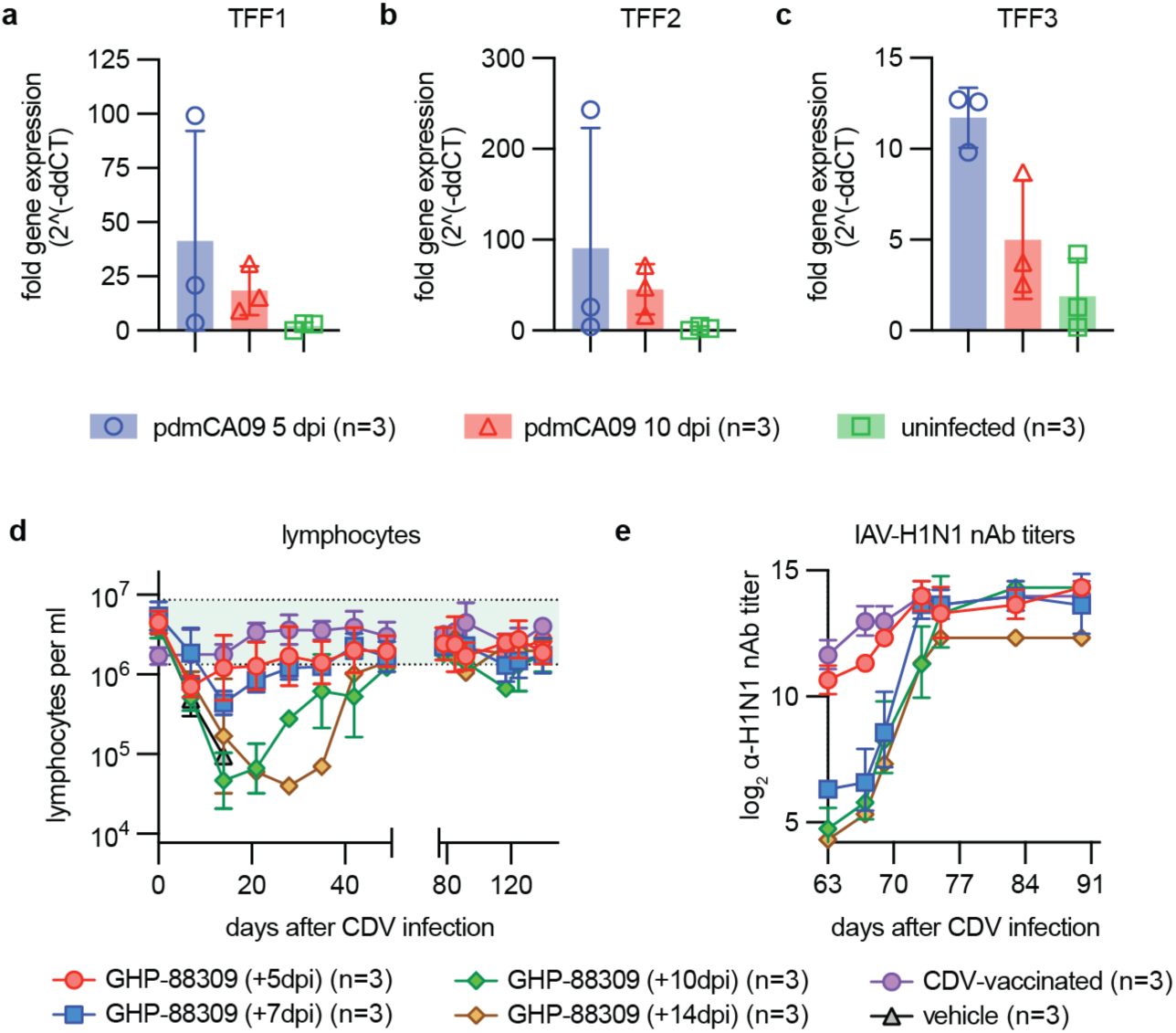
Effect of antiviral treatment on lymphocytopenia and preservation of existing immunity. a-c, RT-qPCR quantitation of relative presence of trefoil factor-encoding message in lung tissue extracted 10 dpi with pdmCA09. **d-e,** Lymphocyte counts (d) and a-pdmCA09 nAb titers (e) from ferrets shown in Fig. 4a; green shading denotes normal range. Symbols represent individual animals, columns show arithmetic means ± SD (a-c), or geometric means ± geometric SD, lines intersect means (d, e); n=3.

