## Supplementary Information for "Therapeutic mitigation of measles-like immune amnesia and exacerbated disease after prior respiratory virus infections in ferrets"

**Supplementary Information table of contents:**

1) Detailed experimental methods
2) Supplementary Table S1. GHP-88309 single oral dose plasma PK in ferrets
3) Supplementary Table S2. GHP-88309 repeat oral dose plasma PK in ferrets
4) Supplementary Table S3. DNA primers used in this study
5) Supplementary Figure S1. Non-formal tolerability study with GHP-88309
6) Supplementary Figure S2. GHP-88309 repeat dose plasma PK
7) Supplementary Figure S3. α-RABV nAbs titers in ferrets before study start.
8) Supplementary Figure S4. Clinical disease after infection of ferrets with recCDV-5804p NΔ425-479 or recCDV-5804p Nectin 4-blind
9) Supplementary Figure S5. CBC results after infection of ferrets with recCDV-5804p NΔ425-479 or recCDV-5804p Nectin 4-blind
10) Supplementary Figure S6. Selected cytokines profiles after flagellin-stimulation of ferrets recovered from CDV
11) Supplementary Figure S7. Challenge of GHP-88309-treated ferrets with CDV-5804p after recovery
12) Supplementary Figure S8. Clinical disease after infection of ferrets with pdmCA09
13) Supplementary Figure S9. α-H1N1 nAbs titers in ferrets before infection with CDV
14) Supplementary Figure S10. Necropsy of consecutively infected ferrets presenting moribund
15) Supplementary Figure S11. All histopathology analyses
16) Supplementary Figure S12. All Gram stains
17) Supplementary Figure S13. Selected cytokines profiles after IAV and CDV infection of ferrets
18) Supplementary Figure S14. Expression of Muc5 proteins in singly or consecutively infected ferrets
19) Supplementary Figure S15. Shed pdmCA09 titers after primary infection of ferrets

20) Supplementary Figure S16. CBC results of consecutively infected ferrets treated with GHP-88309
21) Supplementary Figure S17. α-CDV nAbs titers in GHP-88309 experienced or inexperienced ferrets after recovery from CDV
22) Supplementary References
23) Supplementary Movie S1. Pulmonary ferret MRI of an uninfected animal
24) Supplementary Movie S2. Axial-slice raw data for reconstruction shown in Supplementary Movie S1
25) Supplementary Movie S3. MRI of hemorrhagic pneumonia after consecutive infection with IAV and CDV
26) Supplementary Movie S4. Axial-slice raw data for reconstruction shown in Supplementary Movie S3
27) Supplementary Movie S5. MRI 15 dpi with CDV of consecutively infected ferrets treated with GHP-88309
28) Supplementary Movie S6. Axial-slice raw data for reconstruction shown in Supplementary Movie S5
29) Supplementary Dataset S1. CDV whole genome sequencing analysis after mono-infection
30) Supplementary Dataset S2. Metagenomics analyses
31) Supplementary Dataset S3. CDV whole genome sequencing analysis after consecutive infection
32) Supplementary Dataset S4. All quantitative source data
33) Supplementary Dataset S5. All statistical analyses

### Detailed Methods

#### Cell lines and transfections

MDCK cells (American Type Culture Collection, CCL-34), Human carcinoma (HEp-2, American Type Culture Collection, CCL-23), and African green monkey kidney epithelial cells (CCK-81; canine signaling lymphocytic activation molecule (Vero-cSLAM) were maintained at 37°C and 5% $CO_2$ in Dulbecco's modified Eagle's medium (DMEM) supplemented with 7.5% fetal bovine serum (FBS). All immortalized cell lines used in this study were regularly tested for microbial contamination (6-month intervals).

#### Viruses

recCDV-5804p, recCDV-5804p-Nectin-4-blind-eGFP, and recCDV-5804p-NΔ425-479 stocks were propagated on Vero-cSLAM and titrated by $TCID_{50}$ assay. recRSV-A2-L19 stocks were propagated on HEp-2 cells and titrated by $TCID_{50}$ assay. A/California/7/2009 (H1N1) and A/Wisconsin/67/2005 (H3N2) were propagated on MDCK cells for 2 to 3 days at 37°C. Influenza viruses were titrated by $TCID_{50}$-hemagglutination ($TCID_{50}$-HA) assay on MDCK cells as previously described[1].

**Viral Whole Genome Sequencing**

Whole genome sequencing of recCDV-5804p-NΔ425-479 stocks and from ferret nasal lavage or BALF was performed essentially as previously described[2]. Before cDNA synthesis, genomic DNA was depleted from the RNA extracts with Turbo DNAse (Thermo #AM1907). First-strand cDNA synthesis was performed using SuperScript IV (Thermo) and random hexamers (Thermo) followed by second strand synthesis with Sequenase 2.0 kit (Thermo). Double-stranded cDNA was simultaneously fragmented and barcoded using Nextera chemistry (Nextera XT and Nextera Flex). The resulting libraries were pooled at equal concentrations, and average library size was determined using Agilent DNA D1000 Tape Station kit (Agilent). Pooled libraries were sequenced on a Nextseq 500 or Novaseq 6000. Reads were quality and adapter-trimmed with fastp[3]. The 3 leading bases, 3 trailing bases, and bases with mean phred<20 in a 4-bp sliding window were cut from each read. Reads shorter than 45bp and reads in which >40% bases had phred <15 were removed. Consensus genomes for the recCDV-5804p-NΔ425-479 stocks were generated from the trimmed reads using REVICA (https://github.com/greninger-lab/revica) with the Canine distemper virus strain 5804P (GenBank #AY386316.1) as the initial reference. The LAVA pipeline, available at https://github.com/greninger-lab/lava/tree/Rava_Slippage-Patch, was used to interrogate allele frequency changes from the 425.input consensus genome as previously described[2]. Samples with <10,000 mapped reads or <95% coverage breadth were excluded from variant analysis. Interactive HTML plots highlighting viral allele frequencies relative to inoculum are included as (Supplementary Datasets 1 and 3). Sequence reads have been uploaded to the sequence read archive under BioProject PRJNA1004336.

**Virus yield reduction**

For virus yield-based dose–response assays, cells were infected (M.O.I. = 0.01 $TCID_{50}$ units per cell) in a 24-well plate format with recCDV-5804, recCDV-5804p-NΔ425-479, or drug resistant recombinants, in the presence of serial compound dilutions. Cell-associated progeny virus was harvested 48 h after infection. Viral titers were determined through $TCID_{50}$ titration. Four-parameter variable slope regression modeling was used to determine $EC_{50}$ and $EC_{90}$ concentrations.

**Pharmacokinetics studies in ferrets**

Female ferrets (6 to 10 months of age) received from Triple F Farms were rested for 1 week, randomly assigned to study groups, and dosed orally with GHP-88309 dissolved in 1% methylcellulose. Blood was collected from the anterior vena cava and tissue sampling at the specified time points. Two to three animals per group were sampled for PK analyses. Plasma was separated from blood in microvette CB300 EDTA tubes (2,000×g; 5 min; 4 °C)(Sarstedt Inc) and tissue samples were snap frozen and stored at −80°C prior to analysis by LC-MS/MS. The calibration curve range was 10–100000 ng ml−1 in blank plasma for single-dose PK and 10 ng/ml to 50000 ng/ml for all other studies. Quality control samples of 30, 300, 7500, and 50000 ng/mL of blank plasma were run before and after the single-dose PK samples and quality control samples of 30, 750, and 5000 ng/ml were run before and after all other plasma samples. The calibration curve range was 1.00-2000 ng/ml of tissue lysate for blank tissues. Quality control samples of 3, 30, and 600 ng/ml of tissue lysate in blank tissues were analyzed at the beginning of each sample set. Calibration in each matrix showed linearity with $R^2$ values of >0.99.

**RSV and IAV infections of ferrets**

Female ferrets (6–10 months of age) were purchased from Triple F Farms and housed in an ABSL-2 facility. Prior to study start, ferrets were rested for 1 week, then randomly assigned to study groups. Animals infected with RSV-A2-L19 were in a separate ABSL2 room from influenza infected ferrets. For influenza virus and RSV infections, ferrets were anesthetized using dexmedetomidine/ketamine. Anesthetized ferrets were inoculated intranasally with $1\times10^5$ $TCID_{50}$ units of IAV-A/CA/07/2009 (H1N1) or $1\times10^6$ TCID50 units of RSV-A2-L19 in a volume of 300 µl per nare. Clinical signs (bodyweight and temperature) were monitored once daily. Nasal lavages were performed once daily using 1 ml of PBS containing 2× antibiotics-antimycotics (Gibco). Treatment with 4'-FlU through oral gavage was administered using a once daily (q.d.) regimen.

**CDV infections of ferrets**

Female ferrets (6-10 months of age) were received from Triple F Farms and housed in an ABSL-2 facility. Ferrets were rested for one week prior to the start of each study and randomly assigned into groups. Influenza vaccinations (quadrivalent, Flucelvax; Seqirus, Inc.) were administered at specified time points prior to CDV infection. Prior to infection, ferrets were anesthetized with dexmedetomidine/ketamine, and infected intranasally with $2 \times 10^5$ $TCID_{50}$ of recCDV-5804p, recCDV-5804p-Nectin-4-blind-eGFP or recCDV-5804p-NΔ425-479 in 200µl (100µl per nare). Twice daily treatment with GHP-88309 was initiated at specified times after infection (3, 5, 7, 10, or 24 days after infection). GHP-88309 was administered by oral gavage in 3.5ml 1% methylcellulose and flushed with 3.5 ml high-calorie liquid dietary supplement. Control groups were administered equivalent volumes of 1% methylcellulose. Bodyweight and temperature were monitored daily. Additional monitoring for clinical signs and any potential adverse effects was performed daily. Blood was harvested at specified time points. To assess viremia, PBMCs were isolated from blood by Ficoll gradient centrifugation. Viremia titers were determined by coculturing serial dilutions of purified PBMCs with Vero-cSLAM cells and expressed as $TCID_{50}$ per $10^6$ PBMCs. Complete blood count (CBC) analyses were performed using a VetScan HM5 (Abaxis) in accordance with the manufacturer's protocol. Shed viral loads were determined from nasal lavages and titrated by $TCID_{50}$. For studies involving influenza challenges, surviving CDV-infected ferrets were anesthetized with dexmedetomidine/ketamine, and infected intranasally with $1 \times 10^5$ [recIAV-A/California/07/2009 (H1N1)] or $2 \times 10^5$ $TCID_{50}$HA [A/Wisconsin/67/2005 (H3N2)] in 200µl (100µl per nare). Animals were monitored once daily for clinical signs, including loss of bodyweight and fever. Upper respiratory virus load was measured from nasal lavages performed once daily for up to 7 days post influenza infection.

**Systemic interferon and cytokine profiling**

The relative expression of interferon, cytokines, and interferon-stimulated genes (ISGs) was determined by real-time PCR analysis. RNA was isolated from purified PBMCs that were harvested at different time points after infection. Complementary DNA was synthesized by reverse transcription using SuperScript III (Invitrogen) using oligo-dT primers in accordance with the manufacturer's protocol. Real-time PCR was performed using Fast SYBR Green Master Mix (Applied Biosystems) on a Quantstudio 3 real-time PCR system (Applied Biosystems). Values were normalized to glyceraldehyde-3-phosphate dehydrogenase mRNA, analyzed by the comparative threshold cycle ($\Delta\Delta C_T$) method and expressed relative to mock-infected animals. Sequences of primers are shown in Supplementary Table S3.

**Determination of neutralizing antibody titers**

Plasma was heat inactivated and serially diluted (twofold steps) in serum-free DMEM, mixed with 100 $TCID_{50}$ units of recCDV-5804p, recIAV-A/California/07/2009 (H1N1), or VSV-RABV-G and incubated for 1 h at 37°C. Mixtures were transferred to cell monolayers and virus neutralization was measured after 3 d by visualization of syncytia, hemagglutination assay, or GFP-positive infected cells for recCDV-5804p, recIAV-A/California/07/2009 (H1N1), and VSV-RABV-G, respectively. Equal amounts of virus in FBS, DMEM containing 7.5% FBS, and serum-free DMEM served as controls. Each blood sample was tested in two technical repeats.

**Quantitation of CDV N protein-encoding RNA in PBMCs**

CDV N protein-encoding RNA was detected using the a primer-probe set cdv_n_taq_fw, cdv_n_taq_rev, and cdv_n_probe. RT-qPCR reactions were performed using a QuantStudio 3 real-time PCR system and the QuantStudio Design and Analysis package (version 1.5.2). The CDV N primer-probe set was used with Taqman Fast Virus 1-step master mix (Thermo Fisher Scientific) to detect viral RNA. To calculate RNA copy numbers, a standard curve was created using a linearized pTM1-CDV N plasmid of known concentration as template. Samples were normalized to the numbers of input PBMCs.

**Ferret MRI**

All ferrets were imaged on a high-resolution 7T Bruker (70/20) Biospec MRI scanner at Georgia State University using the ParaVision software package (Bruker, Billerica, MA, version 360.3.4). Animals were anesthetized using a combination of dexamethasone and isoflurane. Respiration rates and body temperature were continuously monitored and maintained using a small-animal physiological monitoring system (SA Instruments Inc, Stony Brook, NY and Kent Scientific, Somnosuite Systems, Torrington, CT). Anesthesia was adjusted to maintain a respiration rate of 40-60 breaths per minute. A 112/86 mm circularly polarized transmitter/receiver coil was employed for in-vivo imaging. Following the localizer scan to position the animal in the center of the magnet, two types of MR protocols were used to capture the lungs in coronal and axial orientations. To overcome the limitations imposed by short T2* of lungs and to reduce motion, a 3D Ultra short echo time (3D UTE) was used to acquire high-quality images with the following parameters: MR acquisition parameters include: 4.3 µs RF block pulse; flip angle (α) = 3.9°; 51,360 radial projections; 128 points on free induction decay (FID); field-of-view (FOV) = 58 × 54 × 67 (mm3); image matrix size = 128 × 128 × 128 (voxels3); receiver bandwidth (BW) = 200 kHz; TE = 0.06 (ms); repetition time (TR) = 4.206 (ms) and 2 signal averages. The total acquisition time for an individual ferret UTE scan was ~7 minutes. An additional 2D T1 IG FLASH (intragate fast low angle shot) self-gated MRI was used to collect images along coronal direction with the following parameters: TR/TE = 400 /3 msec, flip angle = 30°, oversampling =10, 30 slices, matrix size 140 x 128, FOV =70 mm x 64 mm. The total acquisition time was 8 min. The images were converted into DICOM/Nifti formats using Bruker PV-360 software and reconstructed and processed using ImageJ (version 2.9.0) and ITK-SNAP (version 3.6.2-alpha).

**Histopathology**

Pathology scoring was performed according to the following scale: for alveoli, bronchiolitis, and pleuritis scores, scoring was based upon distribution: 0 = no lesions, 1 = focal, 2 = multifocal, 3 = multifocal to coalescing, 4 = diffuse; for perivascular cuffing (PVC) score: 1 = 1 layer of leukocytes surrounding most affected vessel, 2 = 2 - 5 layers, 3 = 6 - 10 layers, 4 = more than 10 layers; for vasculitis score: 1 = infiltration of vessel wall by leukocytes, 2 = infiltration and separation of smooth muscle cells by edema, 3 = same changes as 2 with fibrinoid change, 4 = effacement of the vessel wall; for interstitial pneumonia score: 1 = infiltration of alveolar septa by 1 leukocyte layer thickness, 2 = expansion by 2 leukocyte thickness, 3 = 3 leukocytes thick, 4 = 4 leukocytes thick or more. The sum of individual scores represents a total histopathology score, generated for each animal.

**RNAseq**

Total RNA was treated with Turbo DNAse (ThermoFisher) and used as input for the Illumina Stranded Total RNA (Ribo-Zero Plus Microbiome) library preparation kit using the manufacturer's specifications, then sequenced on an Illumina NovaSeq 6000 to obtain approximately 25 million paired end 150 bp reads. Raw reads were quality- and adapter-trimmed with Trimmomatic v0.39[4]. Metagenomic analysis of trimmed read pairs was performed using the CZID pipeline[5-7] and host and CDV-specific proportions determined. Trimmed read pairs were pseudoaligned to the draft ferret transcriptome MusPutFur1.0, INSDC Assembly GCA_000215625.1[8] using Kallisto v0.46[9] and transcripts aggregated by gene. Genes with an average raw expression level less than 1 raw count per sample were filtered prior to analysis. Differential expression between groups of CDV-infected and flu-pretreated CDV-infected ferret samples collected during survival BAL procedures was calculated with the Wald test in DEseq2[10], with an adjusted *p* value significance threshold of 0.01. All analyses were performed using R v4.2.1. Raw sequence reads used for RNAseq analysis have been uploaded to the sequence read archive, BioProject PRJNA1004336.

**Supplementary Tables**

**Supplementary Table S1. GHP-88309 single oral dose plasma PK in ferrets.** Selected plasma PK parameters of GHP-88309 in ferrets after a single oral dose of 50 or 150 mg/kg.

| **ID** | **dose[A]** [mg/kg] | $\mathbf{t_{max}}$ [hours] | $\mathbf{C_{max}}$ [nmol/ml] | **AUC-INF** [hours×nmol/ml] | **AUC-INF/dose** [hours×nmol/ml/mmol] | $\mathbf{t_{1/2}}$ [hours] |
|---|---|---|---|---|---|---|
| GHP-88309 | 50 | 2 | 55.1 ± 27.5 | 177.8 ± 71.5 | 947.3 ± 381 | 1.5 ± 0.3 |
| GHP-88309 | 150 | 3.3 ± 2.3 | 92.8 ± 30.6 | 754.1 ± 731.7 | 4017.5 ± 3898.4 | 2 ± 0.29 |

[A]Data analysis with WinNonLin; n=3.

**Supplementary Table S2. GHP-88309 repeat oral dose plasma PK in ferrets.** Selected plasma PK parameters of GHP-88309 in ferrets after a single oral dose of 50 or 150 mg/kg

| **ID** | **dose[A]** [mg/kg] | $\mathbf{t_{max}}$ [hours] | $\mathbf{C_{max}}$ [nmol/ml] | **AUC-INF** [hours×nmol/ml] | **AUC-INF/dose** [hours×nmol/ml/mmol] | $\mathbf{t_{1/2}}$ [hours] |
|---|---|---|---|---|---|---|
| GHP-88309 | 15 | 2 | 13.1 ± 0.29 | 49.7 ± 13.7 | 887.1 ± 244.1 | 0.84 ± 0.16 |
| GHP-88309 | 50 | 3 ± 1 | 72.7 ± 17.8 | 359.1 ± 144.9 | 1913.1 ± 772.2 | 1.3 ± 0.66 |

[A]Data analysis with WinNonLin; n=3.

**Supplementary Table S3. DNA primers used in this study.**

| species | target | sequence |
|---|---|---|
| Ferret | il-8_fw | 5'-tgctttctgcagttctgtgtgagc-3' |
| Ferret | il-8_rv | 5'-atgtgggccactgtcaatcactct-3' |
| Ferret | ifn-β_fw | 5'-ggtgtatcctccaaactgctctcc-3' |
| Ferret | ifn-β_rv | 5'-cactccacactgctgctgcttag-3' |
| Ferret | il6_fw | 5'-agtggctgaaacacgtaacaattc-3' |
| Ferret | il6_rv | 5'-atggccctcaggctgaact-3' |
| Ferret | tff1_fw | 5'-ccaaggtggtctgtgttctc-3' |
| Ferret | tff1_rv | 5'-tcctcgtcaggagagttgt-3' |
| Ferret | tff2_fw | 5'-gagcagtgtgtgatggaagt-3' |
| Ferret | tff2_rv | 5'-agatgaaggaaagccaggaag-3' |
| Ferret | tff3_fw | 5'-atgcatcttctcggctgtc-3' |
| Ferret | tff3_rv | 5'-ccactgcacattgcttcaaa-3' |
| Ferret | tgfbeta_fw | 5'-gacatcaacgggctcagttc-3' |
| Ferret | tgfbeta_rv | 5'-gatccacttccagcccagat-3' |
| Ferret | gapdh_fw | 5'-aacatcatccctgcttccactggt-3' |
| Ferret | gapdh_rv | 5'-tgttgaagtcgcaggagacaacct-3' |
| Ferret | ifnγ_fw | 5'-tcaaagtgatgaatgatctctcacc-3' |
| Ferret | ifnγ_rv | 5'-gccgggaaacacactgtgac-3' |
| Ferret | isg15_fw | 5'-agcagcagatagccctgaaa-3' |
| Ferret | isg15_rv | 5'-cagttcttcaccaccagcag-3' |
| Ferret | il1b_fw | 5'-ttctttgaggctgatggtcc-3' |
| Ferret | il1b_rv | 5'-acacgaaatggctcagactc-3' |
| Ferret | tnfa_fw | 5'-ccagatggcctccaactaatca-3' |
| Ferret | tnfa_rv | 5'- ggcttgtcacttggagttcga-3' |
| Ferret | muc5ac_fw | 5'-gcagtccctccaagaatgaa-3' |
| Ferret | muc5ac_rv | 5'-cacacacactggcactgata-3' |
| Ferret | muc5b_fw | 5'-aaacgtcatcgggagtcattag-3' |
| Ferret | muc5b_rv | 5'-atctgggtggtggagatagt-3' |
| CDV | cdv_n_taq_fw | 5'-cgggcaagaaatggtcagaa-3' |
| CDV | cdv_n_taq_rev | 5'-ctgagcctcttccttggtga-3' |
| CDV | cdv_n_probe | fam 5'-acttgccgccgagcttggca-3' bhq-1 |

**Supplementary Figures**

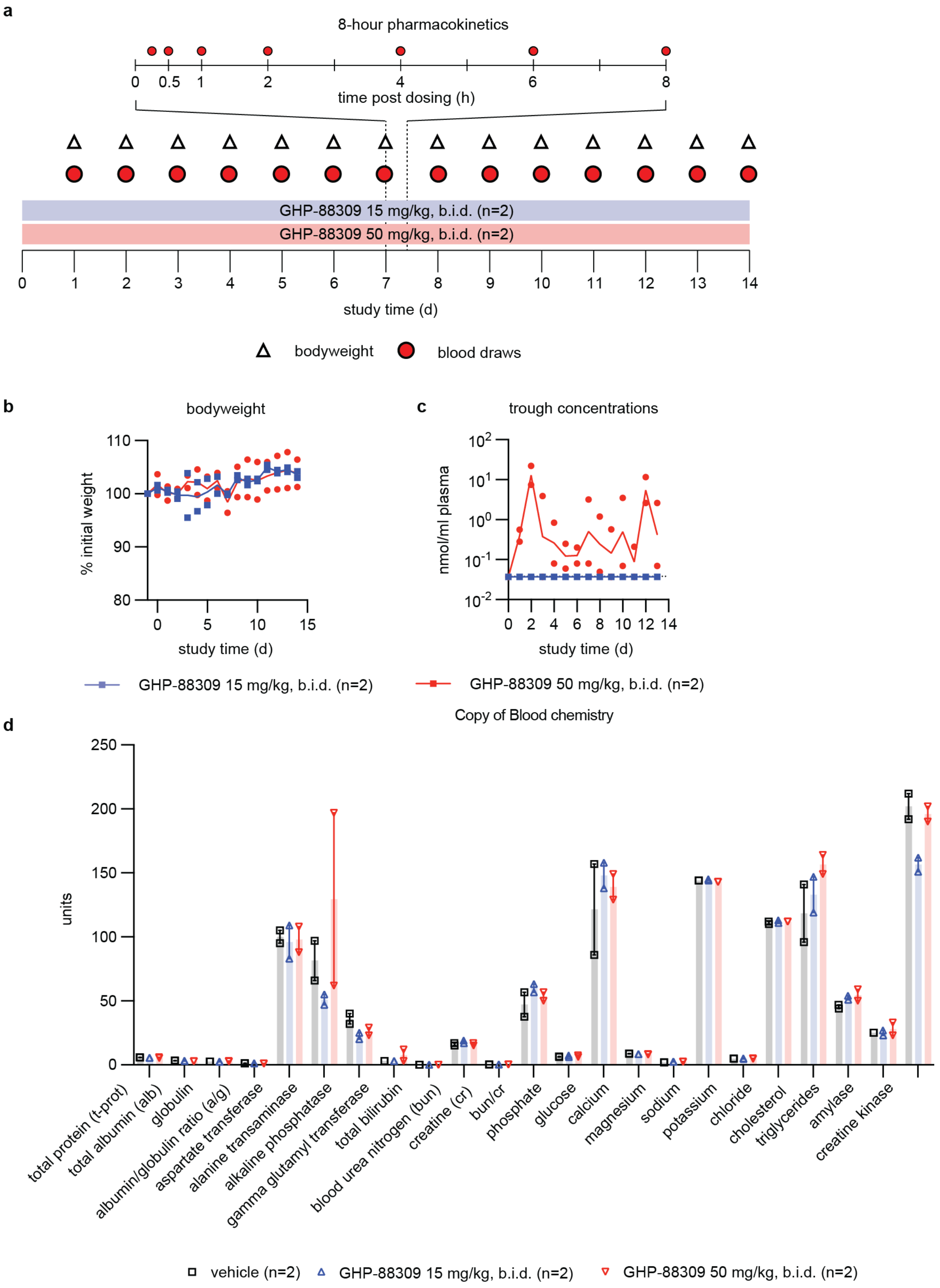


**Supplementary Figure S1. Non-formal tolerability study with GHP-88309. a,** Schematic of the 14-day study. Compound was delivered orally q.d. at 15 and 50 mg/kg. **b-c,** Once daily assessment of bodyweight (b) and GHP-88309 trough plasma concentrations (c). Symbols show averages of 2 animals. **d,** Select blood chemistry parameters. Symbols represent individual animals, columns show averages and range.

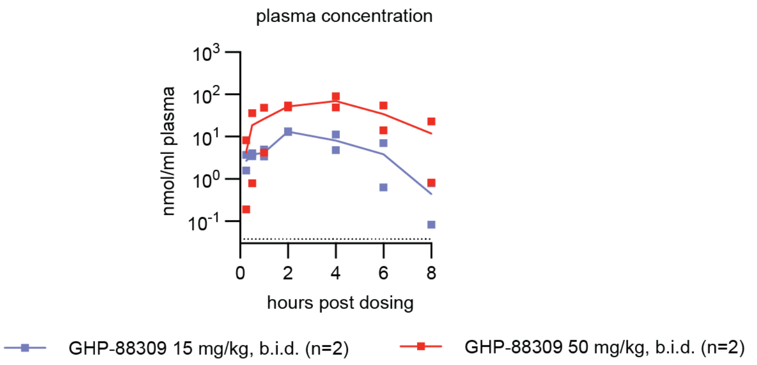


**Supplementary Figure S2. GHP-88309 repeat-dose plasma PK.** Ferrets were orally dosed with GHP-88309 for 7 days q.d. Shown are GHP-88309 plasma concentrations on the last day of dosing. Symbols represent individual animals, lines connect data averages.

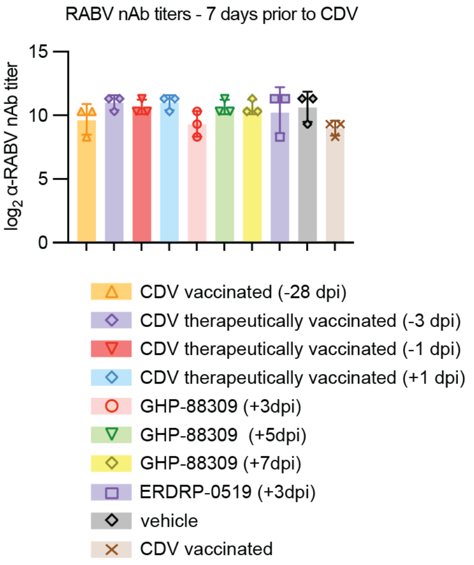


**Supplementary Figure S3. α-RABV nAbs titers in ferrets before study start.** Shown are neutralizing titers determined using a VSV-ΔG pseudotyped with RABV G virus. Symbols represent individual animals, columns represent geometric means ± geometric SD; n=3.

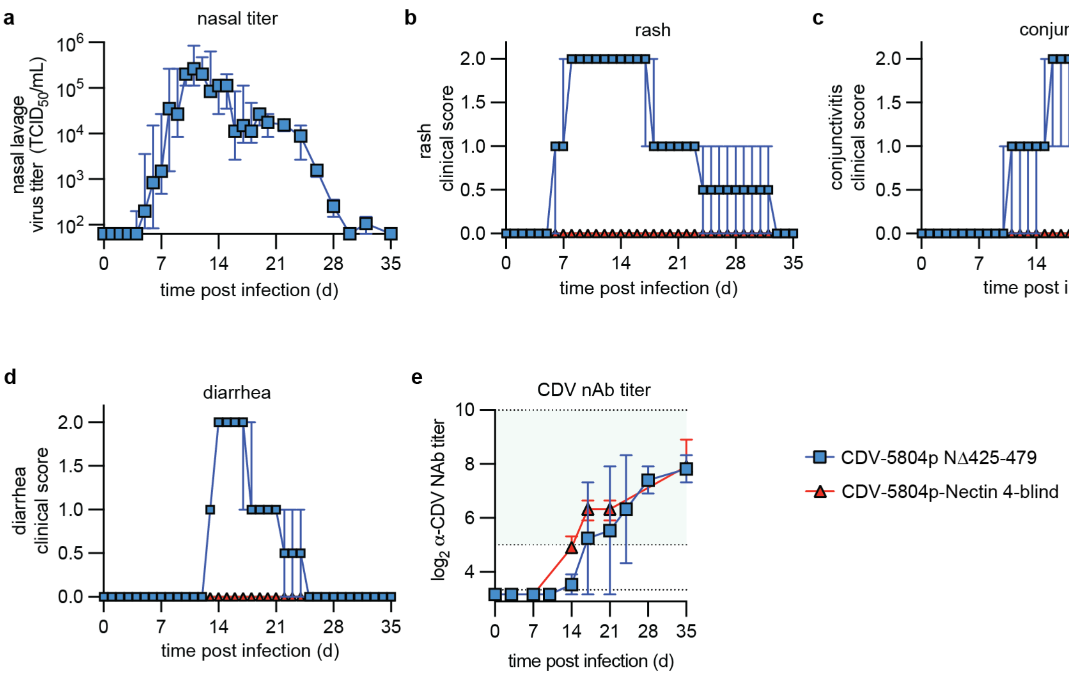


**Supplementary Figure S4. Clinical disease after infection of ferrets with recCDV-5804p NΔ425-479 or recCDV-5804p Nectin 4-blind. a,** Shed virus titer in nasal lavages obtained daily. **b-d,** Presentation of hallmarks of morbillivirus disease including rash (b), conjunctivitis (c), and diarrhea (d). **e,** Appearance of α-CDV nAbs titers in ferrets after infection. Shown are neutralizing titers determined using non-modified recCDV-5804p. Symbols represent geometric means ± geometric SD (a, e) or arithmetic means ± SD; green shading denotes protective nAb titers; n=3.

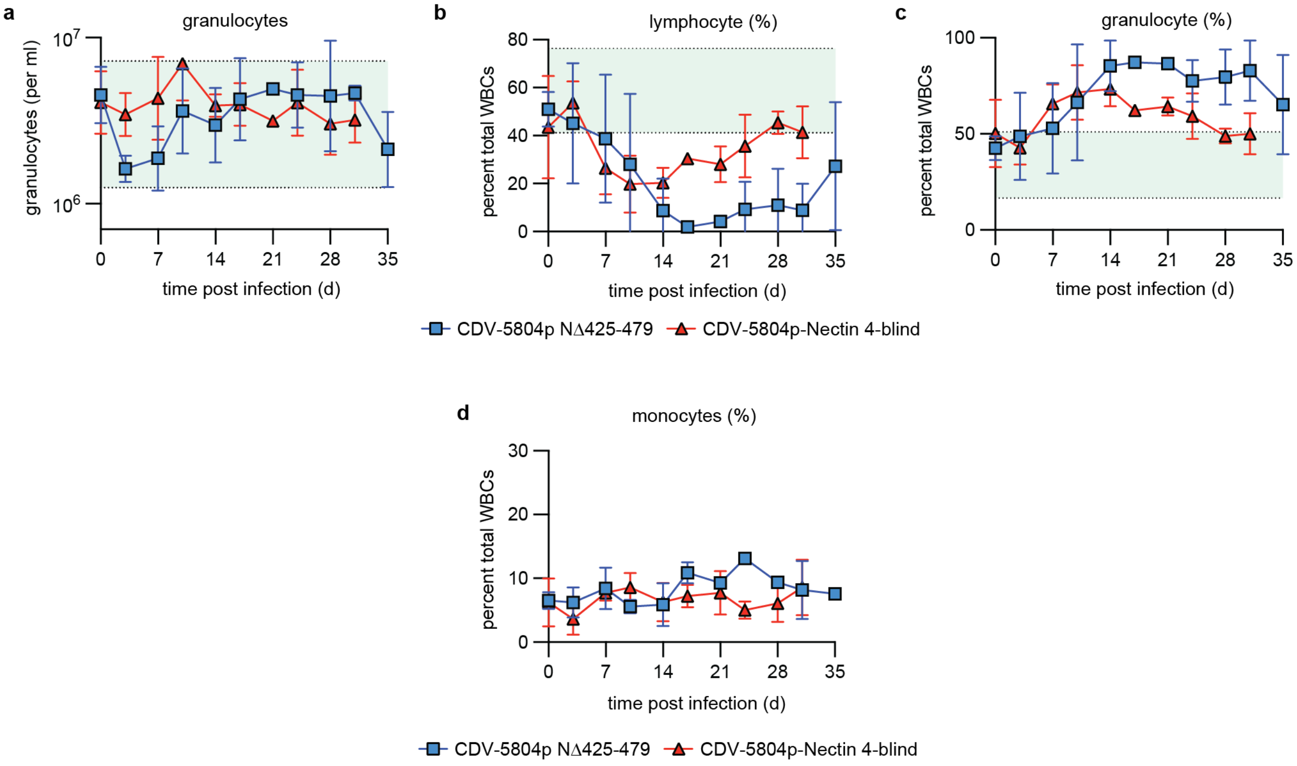


**Supplementary Figure S5. CBC results after infection of ferrets with recCDV-5804p NΔ425-479 or recCDV-5804p**

**Nectin 4-blind. a-d,** Absolute quantitation of neutrophils (a) and relative quantitation of lymphocytes (b), granulocytes (c), and monocytes (d) after CDV infection. Symbols represent geometric means ± geometric SD (a, d) or arithmetic means ± SD (b, c), lines connect means. Green shadings denote normal range in uninfected animals; n=3.

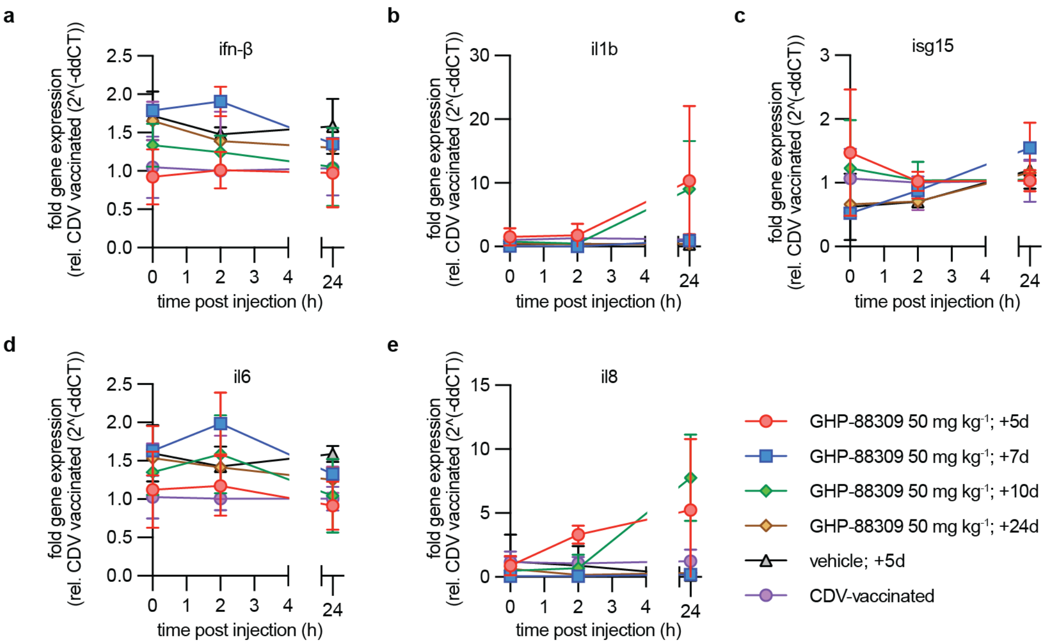


**Supplementary Figure S6. Selected cytokines profiles after flagellin-stimulation of ferrets recovered from CDV. a-e,** Relative changes in expression level of IFN-β (a), IL-1b (b), ISG-15 (c), IL-6 (d), and IL-8 (e) were determined in a 24-hour period after i.m. administration of purified flagellin by RT-qPCR. Symbols represent arithmetic means ± SD, lines connect means; n=3.

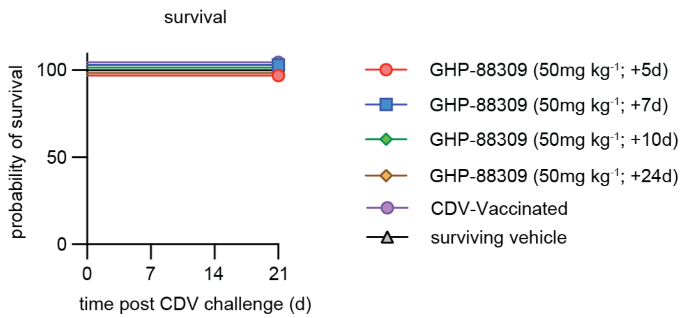


**Supplementary Figure S7. Challenge of GHP-88309-treated ferrets with CDV-5804p after recovery.** Animals were monitored for 21 days after challenge; n=3.

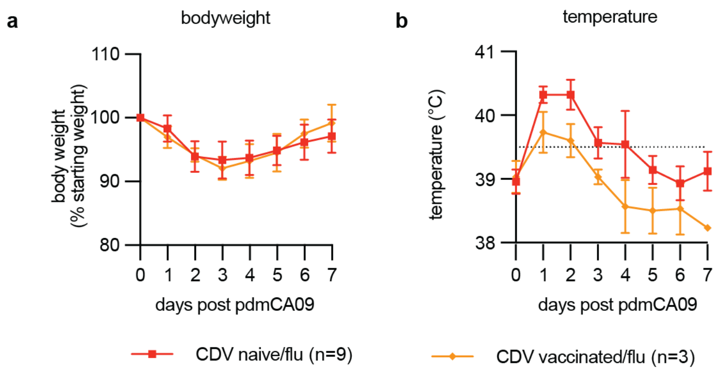


**Supplementary Figure S8. Clinical disease after infection of ferrets with pdmCA09. a-b,** Presentation of hallmarks of IAV disease including loss of bodyweight (a) and fever (b). Symbols represent arithmetic means ± SD, lines connect means; dotted line (b) defines fever in ferrets; n=3 or 9 as specified.

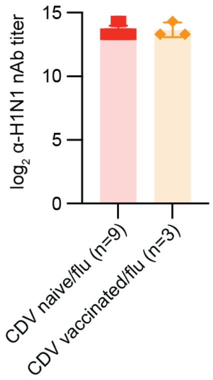


**Supplementary Figure S9. α-H1N1 nAbs titers in ferrets before infection with CDV.** Shown are neutralizing titers determined using pdmCA09. Symbols represent geometric means ± geometric SD; n=3 or 9 as specified.

CDV/Flu (CDV D15) (From Fig. 3)

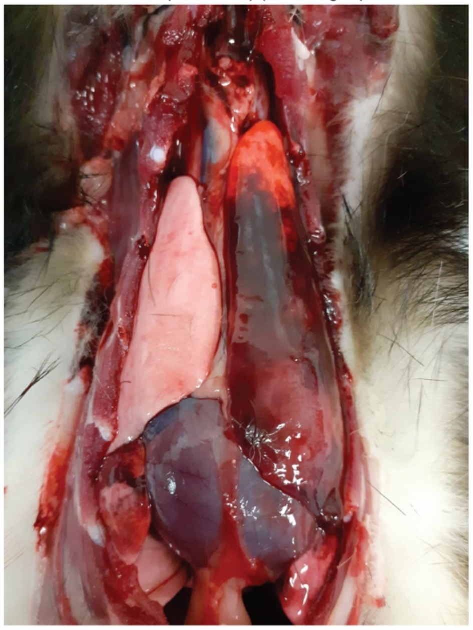

CDV/Flu (CDV D15)

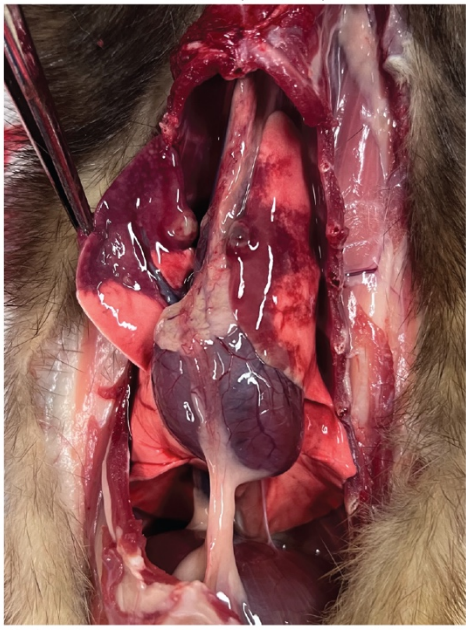

CDV/Flu (CDV D16)

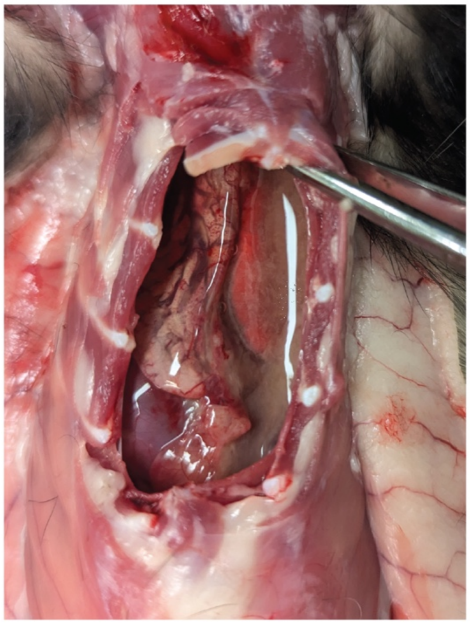

CDV/Flu (CDV D18)

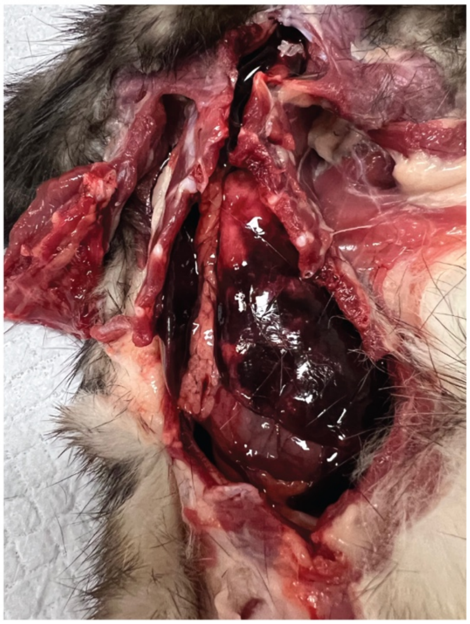

CDV/Flu (CDV D16)

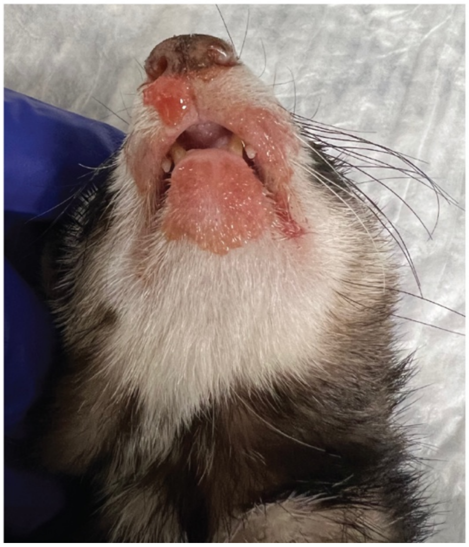

**Supplementary Figure S10. Necropsy of consecutively infected ferrets presenting moribund.** Shown is hemorrhagic pneumonia presentation in independent, consecutively infected animals that could be subjected to necropsy; top left, uncropped image of the animal shown in Fig. 3c.

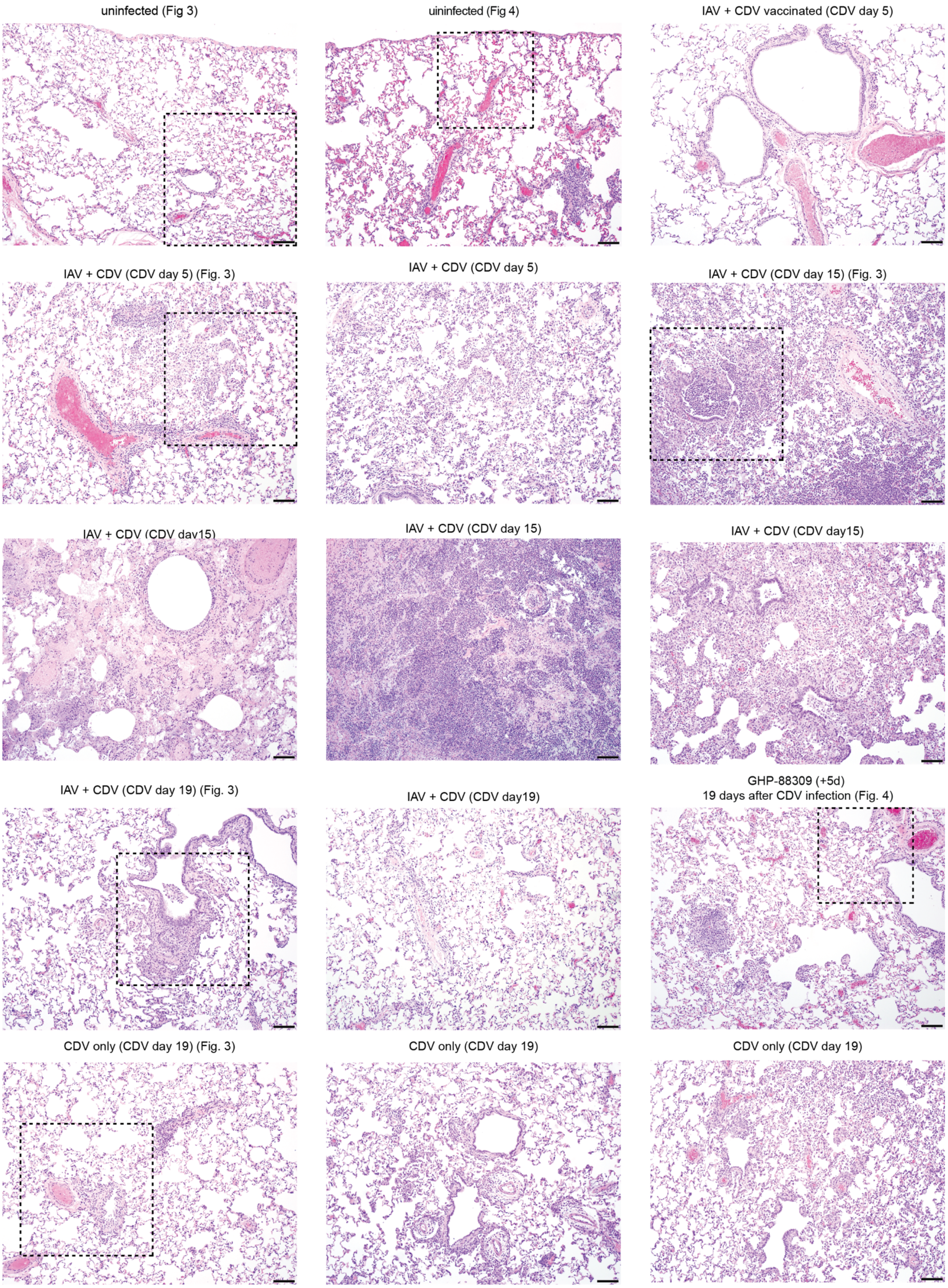
uninfected (Fig 3)
uninfected (Fig 4)
IAV + CDV vaccinated (CDV day 5)
IAV + CDV (CDV day 5) (Fig. 3)
IAV + CDV (CDV day 5)
IAV + CDV (CDV day 15) (Fig. 3)
IAV + CDV (CDV day15)
IAV + CDV (CDV day 15)
IAV + CDV (CDV day15)
IAV + CDV (CDV day 19) (Fig. 3)
IAV + CDV (CDV day19)
GHP-88309 (+5d)
19 days after CDV infection (Fig. 4)
CDV only (CDV day 19) (Fig. 3)
CDV only (CDV day 19)
CDV only (CDV day 19)

**Supplementary Figure S11. All histopathology analyses.** Shown are H&E-stained lung section of independent animals. Dashed squares mark the fields of view shown in Fig. 3d and Fig. 4h, respectively.

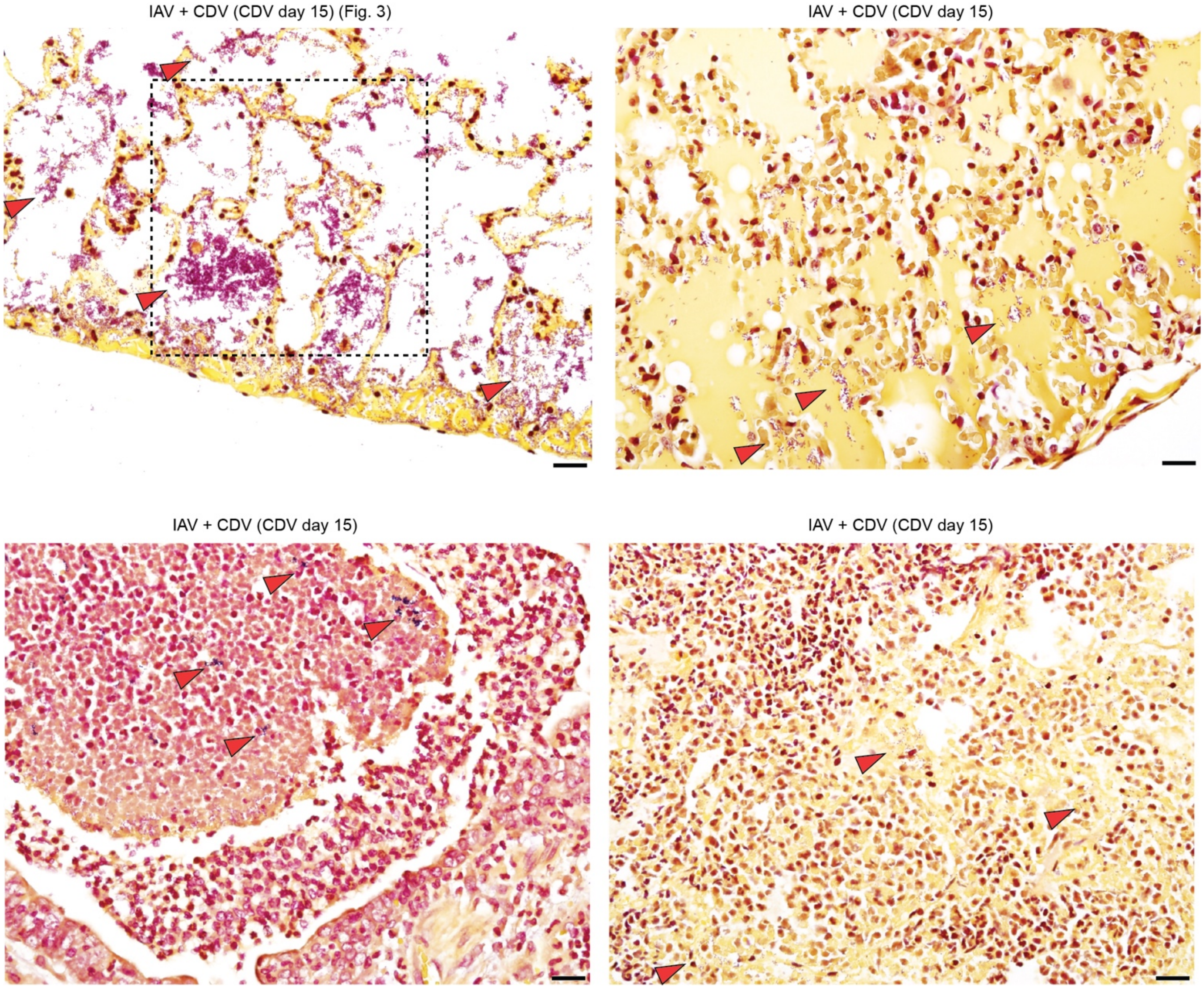


**Supplementary Figure S12. All Gram stains.** Shown are Gram stains of lung sections of independent animals. Dashed square marks the field of view shown in Fig. 3e.

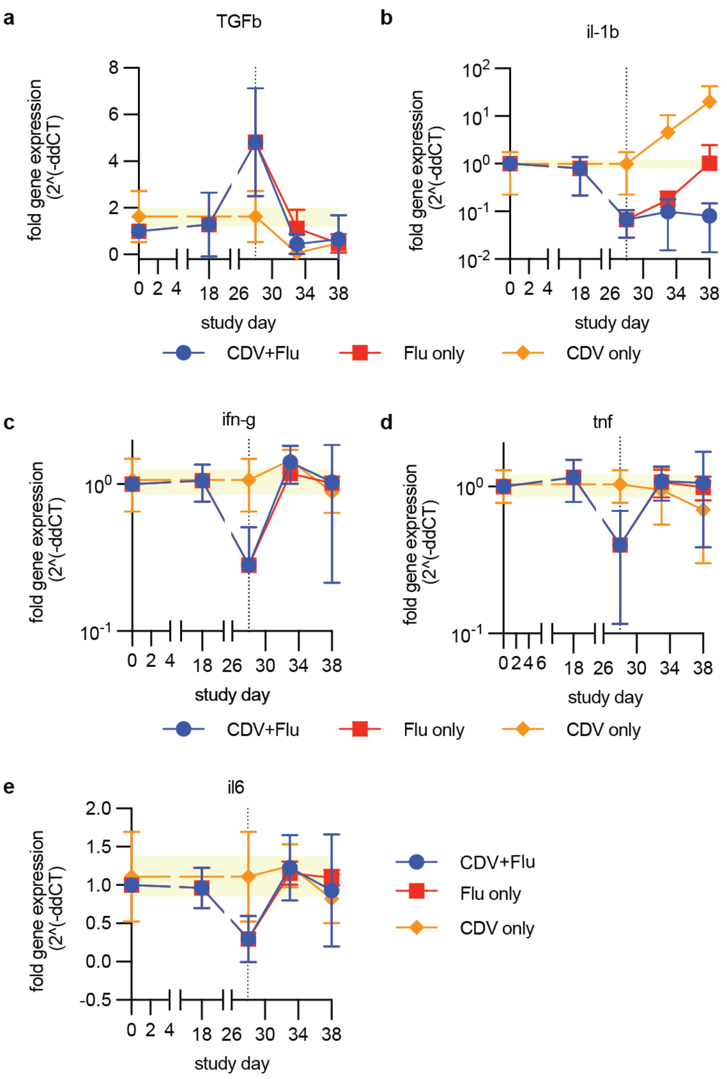


**Supplementary Figure S13. Selected cytokines profiles after IAV and CDV infection of ferrets. a-e,** Relative changes in expression level of TGF-β (a), IL-1b (b), IFN-γ (c), TNF (d), and IL-6 (e) message were determined at the indicated time points by RT-qPCR. Symbols represent arithmetic means ± SD, lines connect means. Dotted lines denote time of infection with CDV; yellow shading specifies variation in uninfected animals; n=3.

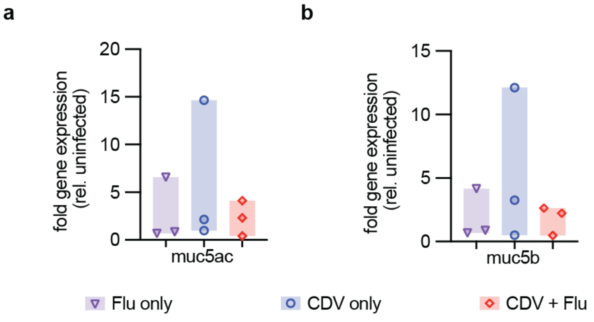


**Supplementary Figure S14. Expression of Muc5 proteins in singly or consecutively infected ferrets. a-b,** Relative changes in expression level of Muc5AC (a) and Muc5B (b) in lung tissues extracted 10 dpi with CDV were determined by RT-qPCR. Symbols represent individual animals, bars denote range from min to max; n=3.

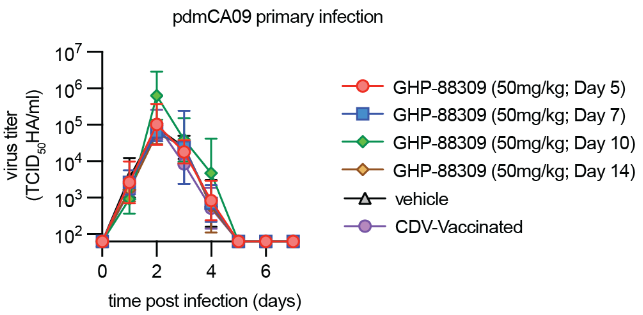


**Supplementary Figure S15. Shed pdmCA09 titers after primary infection of ferrets.** Shed virus titers were determined at the indicated time points through HA-$TCID_{50}$. Symbols represent geometric means ± geometric SD, lines connect means; x-axis intersects at level of detection; n=3.

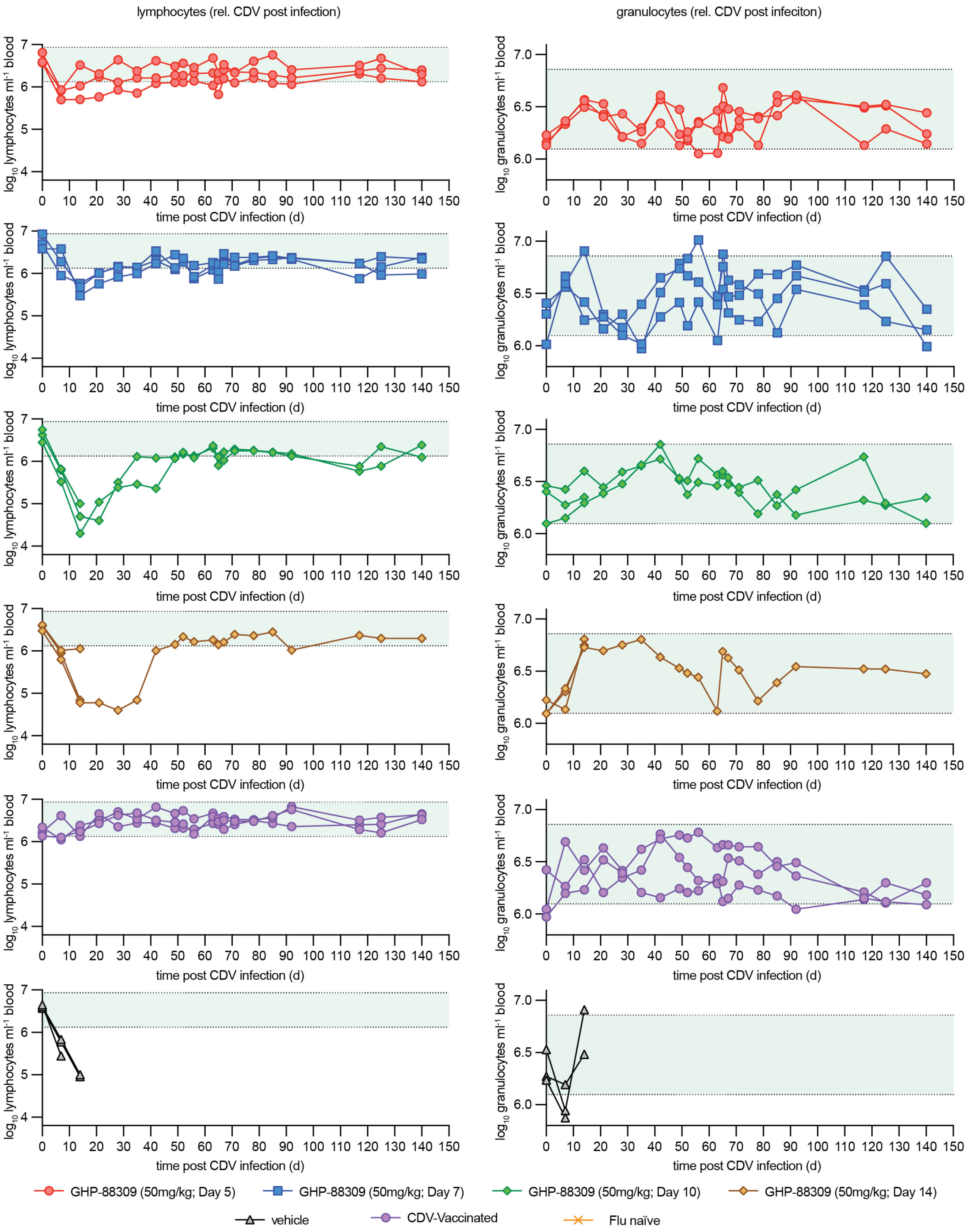


**Supplementary Figure S16. CBC results of consecutively infected ferrets treated with GHP-88309.** Absolute quantitation of lymphocytes (left column) and granulocytes (right column) relative to that before CDV infection. Symbols represent, and lines connect, individual animals. Green shadings denote normal range in uninfected animals; n=3.

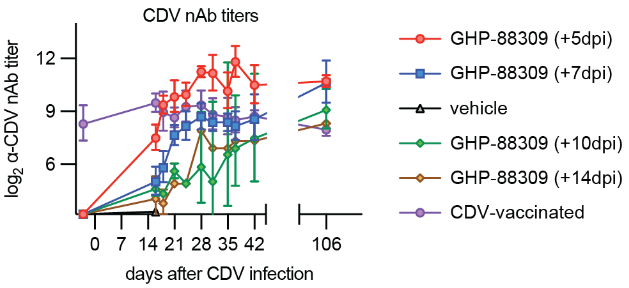


**Supplementary Figure S17. α-CDV nAbs titers in GHP-88309 experienced or inexperienced ferrets after recovery from CDV.** Shown are neutralizing titers determined using recCDV-5804p. Symbols represent geometric means ± geometric SD; n=3.
